# Branched-Chain Amino Acid Metabolic Reprogramming Orchestrates Drug Resistance to EGFR Tyrosine Kinase Inhibitors

**DOI:** 10.1101/638007

**Authors:** Yuetong Wang, Jian Zhang, Shengxiang Ren, Dan Sun, Hsin-Yi Huang, Hua Wang, Yujuan Jin, Fuming Li, Chao Zheng, Liu Yang, Lei Deng, Zhonglin Jiang, Tao Jiang, Xiangkun Han, Shenda Hou, Chenchen Guo, Fei Li, Dong Gao, Jun Qin, Daming Gao, Luonan Chen, Kwok-Kin Wong, Cheng Li, Liang Hu, Caicun Zhou, Hongbin Ji

## Abstract

Drug resistance is a significant hindrance to effective cancer treatment. Although resistance mechanisms of epidermal growth factor receptor (EGFR)-mutant cancer cells to lethal EGFR tyrosine kinase inhibitors (TKI) treatment have been investigated intensively, how cancer cells orchestrate adaptive response under sublethal drug challenge remains largely unknown. Here we find that 2-hour sublethal TKI treatment elicits a transient drug-tolerant state in EGFR-mutant lung cancer cells. Continuous sublethal treatment reinforces this tolerance and eventually establishes long-term TKI resistance. This adaptive process involves H3K9 demethylation-mediated epigenetic upregulation of branched-chain amino acid aminotransferase 1 (BCAT1) and subsequent metabolic reprogramming, which promotes TKI resistance through attenuating reactive oxygen species (ROS) accumulation. Combinational treatment with TKI and ROS-inducing reagents overcomes this drug resistance in preclinical mouse models. Clinical information analyses support the correlation of BCAT1 expression with EGFR TKI response. Collectively, our findings reveal the importance of epigenetically regulated BCAT1-engaged metabolism reprogramming in TKI resistance in lung cancer.

**HIGHLIGHTS:** Sublethal EGFR TKI treatment induces transient drug-tolerant state and long-term resistance in EGFR-mutant lung cancer cells

Epigenetically regulated BCAT1-mediated metabolic reprogramming orchestrates EGFR TKI-induced drug resistance

Combinational treatment with TKI and ROS-inducing agents overcomes the drug resistance induced by EGFR TKI treatment

## INTRODUCTION

Cancer cells are notorious for its strong plasticity in response to cytotoxic stress. They may modulate adaptive programs to survive during therapy (Holohan et al., 2013). Upon lethal EGFR TKI exposure, drug-sensitive cancer cells initially undergo drastic apoptotic cell death resulting in notable tumor regression; however, the remaining drug-tolerant cells can follow distinct evolutionary paths and eventually acquire resistance through mutation or non-mutation mechanisms (Hata et al., 2016). However, little is known about how cancer cells orchestrate their adaptive response under sublethal TKI challenge.

Epigenetic alterations are one of the major mechanisms underlying the adaptation of cancer cells to drug exposure (Easwaran et al., 2014). Epigenetic changes such as H3K4 demethylation or H3K9 and H3K27 methylation has been linked to lethal EGFR TKI induced resistance (Guler et al., 2017; Sharma et al., 2010).

Under stressful conditions, cancer cells can reprogram cellular metabolism to support their malignant phenotypes including proliferation, invasion and resistance to drug therapy (Vander Heiden and DeBerardinis, 2017). Recently, Aberrant branched-chain amino acids (BCAAs) metabolism has recently been implicated in various human malignancies including leukemia, liver cancer, pancreatic cancer and non-small cell lung cancer (Ananieva and Wilkinson, 2018; Ericksen et al., 2019; Hattori et al., 2017; Mayers et al., 2016). In addition, global metabolic changes including BCAAs catabolism alteration is observed in lung adenocarcinoma cells during 9 days of lethal EGFR TKI treatment (Thiagarajan et al., 2016). In particular, growing evidence has highlighted an essential role for BCAT1, a cytosolic aminotransferase that catalyzes the catabolism of BCAAs, in promoting cancer progression (Hattori et al., 2017; Thewes et al., 2017; Tonjes et al., 2013). However, whether BCAT1-mediated metabolic program is involved in EGFR TKI-induced drug resistance remains unclear.

Here, we demonstrate that sublethal EGFR TKI exposure for 2 hours elicits a transient drug-tolerant state in human EGFR-mutant lung cancer cells. Such short-term tolerance could be reinforced by continuous sublethal TKI treatment and eventually establish long-term drug resistance. Mechanistically, we demonstrate that this drug resistance is potentially mediated through BCAT1-mediated metabolic reprogramming via epigenetic regulation.

## RESULTS

### Sublethal EGFR TKI Treatment Induces Transient and Long-Term Resistance

To identify a suitable sublethal EGFR TKI dose, we treated the EGFR-mutant lung cancer cell line PC9 with different drug concentrations and found that 10nM gefitinib (GEF) treatment effectively blocked the activation of EGFR without inducing a significant cell apoptosis, whereas drug concentration over 100nM triggered overt apoptosis (Figure 1A-B and S1A). Thus we employed this sublethal dose for further study. We performed the short-term sublethal treatment experiments: cancer cells were exposed to sublethal TKI for 2 hours (hrs) followed by recovery in drug-free medium for different time intervals before re-exposed to higher doses of TKI for 0.5 hr; then cells were cultured in drug-free medium for additional 72 hrs before MTT assay (Figure 1C). Notably, sublethal GEF pre-treatment rendered PC9 cells more tolerant to ensued higher doses of GEF treatment (Figure 1D). Similar drug-tolerance state was also observed in other EGFR-mutant lung cancer cell lines HCC827 and SH450 (Zheng et al., 2011) (Figure 1D and S1B). We found the drug-tolerance effect was most significant when cells were re-exposed to 50nM GEF but decreased with increasing drug concentration. Such TKI tolerance state could maintain over 2hrs after drug withdrawal and diminished gradually over time (Figure 1D). This tolerance was delicately regulated as when higher doses of GEF (0.1-1μM) were used for pre-treatment, no significant tolerance was detected (Figure S1C), which might be due to increased cytotoxicity of pre-treatment at such high concentration. Similar drug-tolerant state was also detected when cells were pre-treated with sublethal dose of erlotinib (ERL) (Figure S1D). Consistently, three cycles of TKI pre-treatment (each cycle contains sublethal drug treatment for 2 hrs followed by recovery in drug-free medium for additional 2 hrs) rendered PC9 and HCC827 cells more resistant to subsequent GEF treatment (Figure 1E-F). Remarkably, continuous sublethal GEF treatment for over three months conferred these cell lines with strong resistance to GEF. Hereafter, we referred to these <u>S</u>ublethal <u>T</u>KI <u>A</u>dapted <u>C</u>ells as STACs. e.g., STAC-P and STAC-H were derived from PC9 and HCC827 cells, respectively (Figure 1G-H). The established STACs had significant increased IC50 value compared with parental cells *in vitro* (Figure S1E). Consistently, STAC-P xenograft tumors showed more resistance to GEF treatment than PC9 xenograft counterparts even at two-fold of regular dose (50 mg/kg daily) (Reagan-Shaw et al., 2008), despite comparable inhibition of EGFR phosphorylation (Figure 1I-J and S1F). Similarly, STAC-H xenograft tumors displayed more resistant to GEF than HCC827 xenograft controls (Figure S1G-H). Additionally, STAC exhibited cross-resistance to ERL (Figure 1K). Even drug withdrawal for over 70 passages, STAC still exhibited strong resistance to TKI (Figure 1L), indicating persistent drug resistance. Collectively, these results demonstrate that sublethal TKI treatment enables EGFR-mutant cancer cells to obtain a transient tolerant state through short-term treatment and persistent resistance to EGFR TKI through long-term treatment.

**Figure 1.**
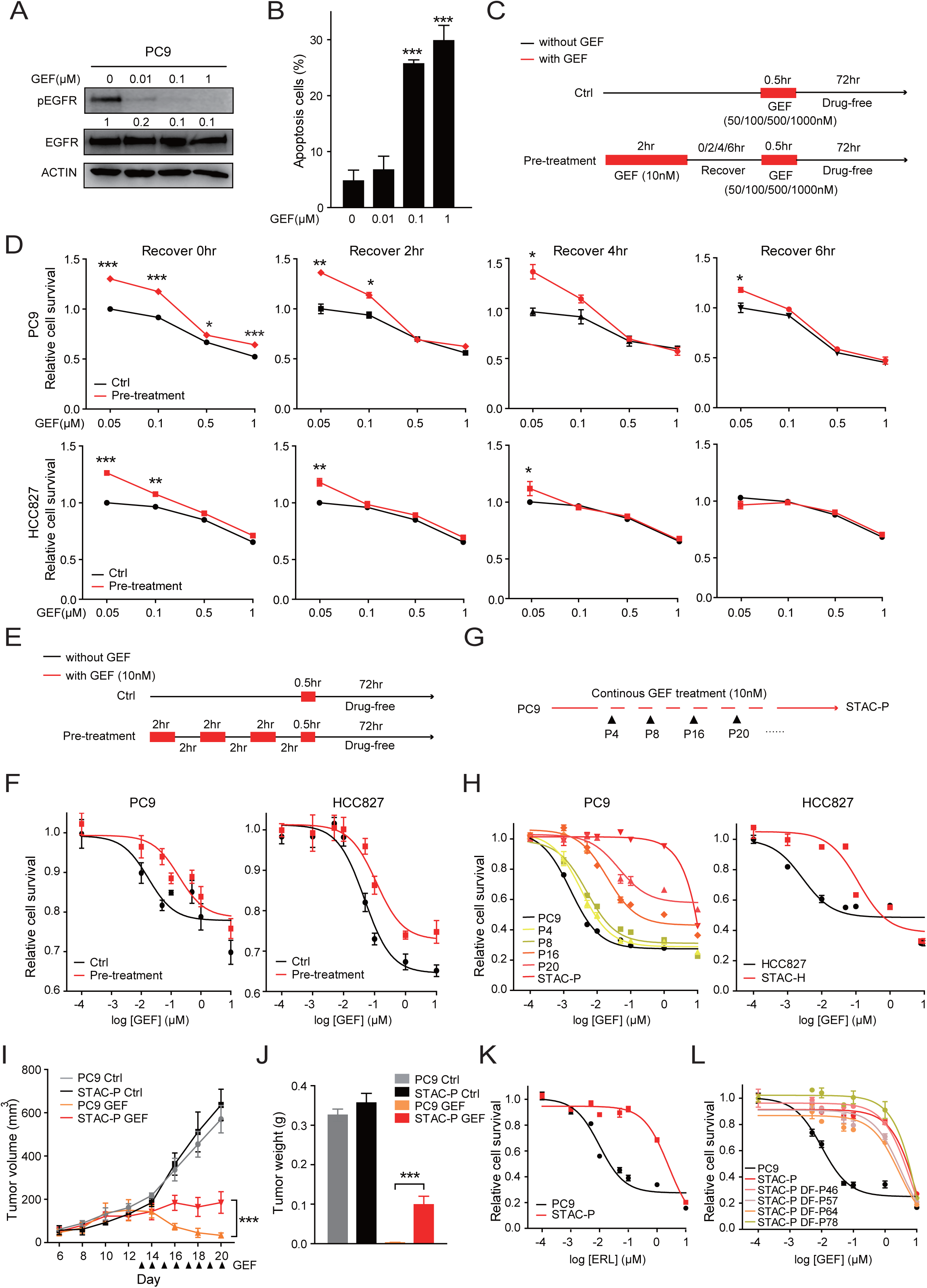
Sublethal EGFR TKI Treatment Induces Short-Term Tolerance and Long-Term Resistance In Human EGFR-Mutant Lung Cancer Cells. (A) PC9 cells were treated with different doses of GEF for 2 hrs and protein levels of pEGFR and total-EGFR were determined by western blot assay. β-actin was used as a loading control. (B) Apoptosis of PC9 cells treated with indicated doses of GEF for 24 hrs by Annexin-V staining. (C) Scheme for short-term sublethal GEF treatment experiments. (D) Cells were exposed to sublethal GEF (10nM) for 2 hrs followed by recovery in drug-free medium with different times before re-exposed to indicated doses of GEF for 0.5 hr, and then cultured in drug-free medium for additional 72 hrs before MTT assay. (E) Scheme for three cycles of short-term sublethal GEF treatment experiments. Each cycle contains exposure to sublethal GEF (10nM) for 2 hrs followed by recovery in drug-free medium for 2 hrs. (F) Cells were exposed to three cycles of short-term sublethal GEF treatment before re-exposed to indicated doses of GEF for 0.5 hr, and then cultured in drug-free medium for additional 72 hrs before MTT assay. (G) Scheme for the establishment of drug-resistant PC9 cells (STAC-P) using continuous sublethal GEF (10nM) treatment for over three months. (H) MTT assay of different passages of PC9 cells continuously exposed to sublethal GEF treated with indicated doses of GEF for 72 hrs (left). MTT assay of HCC827 and STAC-H cells treated with indicated doses of GEF for 72 hrs (right). (I) Growth curves of xenograft tumors derived from PC9 and STAC-P cells treated with or without GEF (50mg/kg) daily via intraperitoneal injection. (n=6 per group) (J) Tumor weight for xenografts treated as in Figure 1I. Data were presented as mean ± SEM. (K) MTT assay of PC9 and STAC-P cells treated with indicated doses of erlotinib (ERL) for 72 hrs. (L) MTT assay of STAC-P cells after GEF withdrawal for different passages treated with indicated doses of GEF for 72 hrs. DF, drug-free. \**p*<0.05, \*\**p*<0.01, \*\*\**p*<0.001.

### Decreased H3K9 methylation Is Involved in TKI Resistance in STAC

Next, we sought to examine the underlying mechanism responsible for EGFR TKI resistance in STAC. Several mechanisms have been proposed for TKI resistance (Rotow and Bivona, 2017). Nevertheless, neither EGFR T790M nor KRAS mutation was detected in STAC using amplification-refractory mutation system (ARMS) (Figure S2A). In addition, STAC and parental cells showed similar proliferation rate and cell cycle distribution (Figure S2B-C). Although phospho-RTK array identified several components of RTK pathways upregulated in STAC (Figure S2D), knockdown of these molecules individually failed to abrogate drug resistance of STAC (Figure S2E-F), indicating other mechanisms.

Previous studies reveal the importance of dysregulation of histone H3 methylation in TKI resistance (Guler et al., 2017; Sharma et al., 2010). We therefore evaluated a series of histone H3 methylation patterns in these TKI-resistance cells. Among the detected H3 methylation, levels of H3K9me2 and H3K9me3 decreased most significantly in STACs (Figure 2A and S3A). Comparable decrease of H3K9me2 and H3K9me3 levels indicates that decreased methylation might occur at the step from H3K9me1 to H3K9me2. Histone methylation is dynamically controlled by methyltransferases and demethylases. We found that the activity of H3K9 methyltransferase, rather than its demethylase activity, was reduced in STACs (Figure 2B and S3B). Consistently, no significant changes of the expression of H3K9 demethylases including JMJD family members (Lim et al., 2010) were observed between STAC and parental cells (Figure S3C), and knockdown of these demethylases individually failed to reverse STAC TKI resistance (Figure S3D-E). Notably, although no substantial differences in protein levels of G9a or SUV39H1, two known H3K9 methyltransferases (Shinkai and Tachibana, 2011; Wang et al., 2012), were observed between STACs and parental counterparts (Figure S3F), knockdown of either enzyme conferred PC9 cells resistance to GEF without notable impact upon cell proliferation (Figure 2C-D and S3G). Similar results were also observed in HCC827 cells (Figure S3H-I). In addition, ectopic expression of shRNA-resistant G9a or SUV39H1 rendered G9a- or SUV39H1-knockdowned cells regained GEF sensitivity, respectively (Figure 2C-D and S3H-I), confirming that the gene knockdown effect upon drug response are on target. Similarly, treatment of PC9 or HCC827 cells with BIX01294, the H3K9 methyltransferase inhibitor (Janzen et al., 2010), promoted GEF resistance (Figure 2E and S3J). Conversely, ectopic expression of G9a or SUV39H1 alone partially restored GEF sensitivity in STACs (Figure 2F-G and S3K). Since DNA demethylation can also be regulated by the DNA demethylase TET (Wu and Zhang, 2017), we then evaluated the enzyme activity of TET by evaluating the levels of 5mC and 5hmC. As shown in Figure S4A-B, the levels of 5mC were unchanged between PC9 vs. STAC-P and HCC827 vs. STAC-H. Interestingly, the levels of 5hmC were decreased in STAC-P and STAC-H when compared to their corresponding controls, indicating that TET activity is reduced in these drug-resistant cells. However, knockdown of TET1, TET2, or TET3 by shRNA had no major effect on GEF sensitivity in either parental cell lines or their STAC-derivatives (Figure S4C-G), indicating that TET may not be involved in mediating TKI resistance in this setting. Thus, these data reveal that STAC TKI resistance involves H3K9 demethylation.

**Figure 2.**
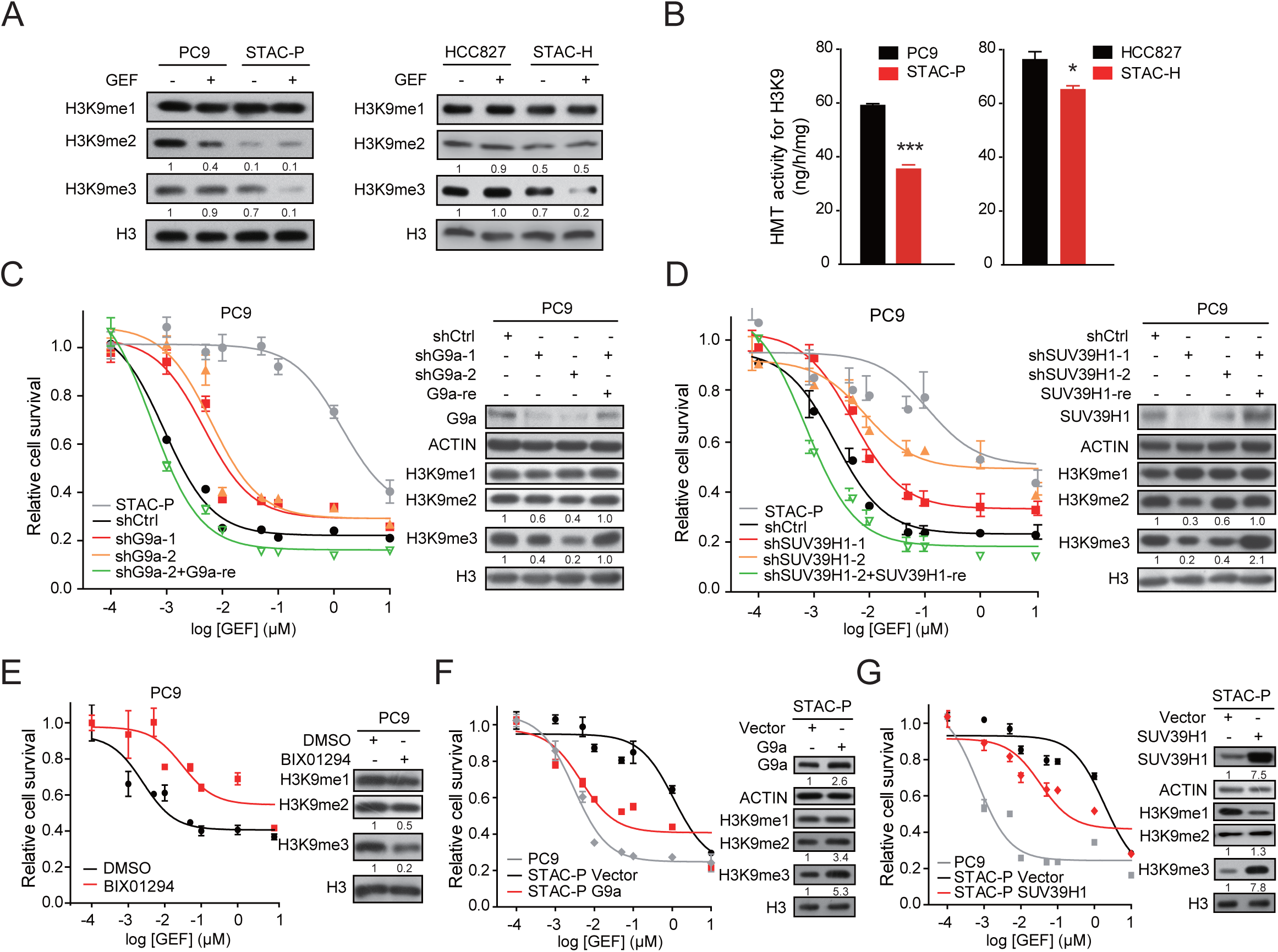
Decreased H3K9 Methylation Is Involved in STAC TKI Resistance. (A) Cells were treated with or without sublethal GEF (10nM) for 2 hrs and lysates were subjected to western blot assay. Histone H3 and β-actin were used as loading controls. (B) Histone methyltransferase (HMT) activities for H3K9 in indicated cells. (C-D) MTT assay of PC9 cells with G9a knockdown or in combination with ectopic expression of shRNA-resistant G9a (C) or with SUV39H1 knockdown or in combination with ectopic expression of shRNA-resistant SUV39H1 (D) treated with indicated doses of GEF for 72 hrs. (E) MTT assay of PC9 cells treated with DMSO or BIX01294 (1µM) together with indicated doses of GEF for 72 hrs. (F-G) MTT assay of STAC-P cells with or without ectopic expression of G9a (F) or SUV39H1 (G) treated with indicated doses of GEF for 72 hrs. \**p*<0.05, \*\*\**p*<0.001.

### Epigenetically upregulated BCAT1 contributes to TKI resistance in STAC

Demethylation of H3K9 is important for de-repression of target gene transcription (Metzger et al., 2005). We then explored which epigenetically regulated molecular events may be involved in mediating STAC TKI resistance. Integrative analyses of microarray and RNA-seq data revealed 22 genes consistently upregulated in STAC (Figure 3A and S5A). Interestingly, individual knockdown of these genes identified BCAT1 as the top hit in reversing TKI resistance in STAC (Figure 3B and S5B). Indeed, BCAT1 was significantly upregulated in STACs both *in vitro* and *in vivo* (Figure 3C-D and S5C-E). In addition, the expression of BCAT1 increased while level of H3K9me2 decreased gradually during the establishment of STAC (Figure S5F), indicating the negative regulation of BCAT1 expression by H3K9 methylation.

**Figure 3.**
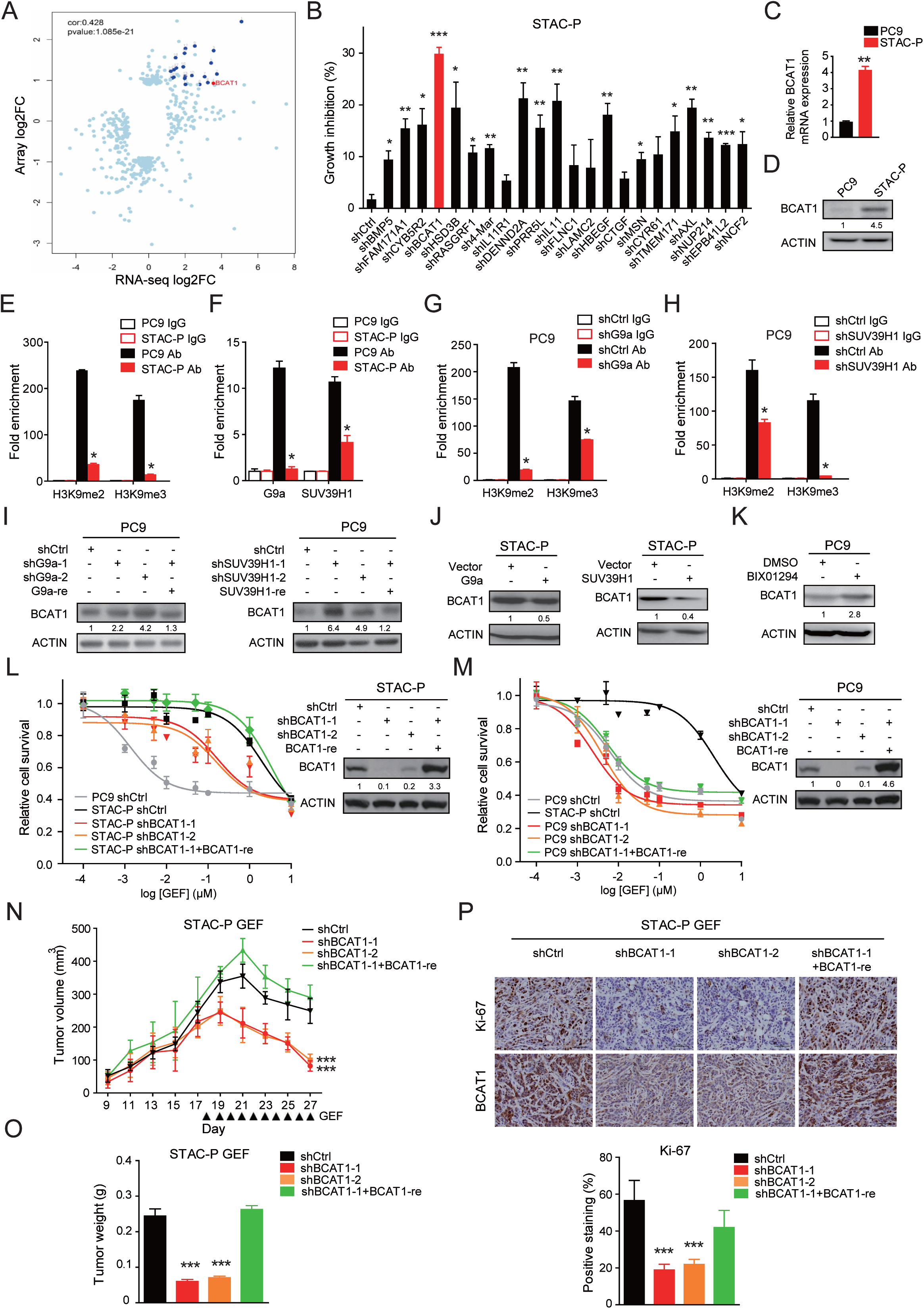
BCAT1 Knockdown Sensitizes STAC to GEF Treatment. (A) Plots showing the significantly dysregulated genes in STAC-P as compared to parental PC9 cells revealed by microarray and RNA-seq analyses. A total of 22 genes were indicated (21 in blue and BCAT1 in red). (B) Growth inhibition rate of STAC-P cells with indicated gene knockdown were determined by MTT assay following GEF treatment (10nM) for 72 hrs. (C-D) Relative mRNA (C) and protein (D) levels of BCAT1 in indicated cells. (E) ChIP assay of the H3K9me2 or H3K9me3 enrichment on *BCAT1* promoter relative to immunoglobulin G (IgG) in indicated cells. (F) ChIP assay of the G9a or SUV39H1 enrichment on *BCAT1* promoter relative to IgG in indicated cells. (G-H) ChIP assay of the H3K9me2 or H3K9me3 enrichment on *BCAT1* promoter relative to IgG in PC9 cells with or without G9a (G) or SUV39H1 (H) knockdown. (I) Protein levels of BCAT1 in PC9 cells with or without G9a or SUV39H1 knockdown. (J) Protein levels of BCAT1 in STAC-P cells with or without ectopic expression of G9a or SUV39H1. (K) Protein levels of BCAT1 in PC9 cells treated with DMSO or BIX01294 (1μM) for 24 hrs. (L-M) MTT assay of STAC-P (L) or PC9 (M) cells with BCAT1 knockdown or in combination with ectopic expression of shRNA-resistant BCAT1 treated with indicated doses of GEF for 72 hrs. (N) Growth curves of xenograft tumors derived from STAC-P cells with BCAT1 knockdown or in combination with ectopic expression of shRNA-resistant BCAT1 following GEF treatment (50mg/kg) daily via intraperitoneal injection. (n=6 per group) (O) Tumor weight for STAC-P xenografts treated as in Figure 3N. Plots were presented as mean ± SEM. (P) Representative immunohistochemical (IHC) staining of Ki-67 in STAC-P xenografts treated as in Figure 3N. Scale bar, 50μm. Statistical analyses of Ki-67 staining were presented as mean ± SEM. \**p*<0.05, \*\**p*<0.01, \*\*\**p*<0.001.

Next we performed chromatin immunoprecipitation (ChIP) assay and found that levels of both H3K9me2 and H3K9me3 at the *BCAT1* promoter were reduced in STACs as compared to parental controls (Figure 3E and S5G). In addition, we confirmed by ChIP that the binding of G9a and SUV39H1 to the *BCAT1* promoter region was also reduced in STACs relative to control cells (Figure 3F and S5H). Moreover, ChIP analysis of H3K9me2 and H3K9me3 in PC9 cells confirmed that knockdown of G9a or SUV39H1 resulted in decreased H3K9me2/3 marks at the *BCAT1* promoter (Figure 3G-H). In parallel, knockdown of G9a or SUV39H1 in PC9 cells upregulated BCAT1 expression; conversely, ectopic expression of either enzyme in STAC-P did the opposite (Figure 3I-J). Similarly, BIX01294 treatment also reduced the H3K9me2/3 marks at the *BCAT1* promoter as revealed by ChIP analysis (Figure S5I), meanwhile increased BCAT1 expression in PC9 cells (Figure 3K). Similar results were also obtained in HCC827/STAC-H cells (Figure S5J-L). These data suggest that upregulation of BCAT1 in STACs is potentially mediated through epigenetic derepression involving H3K9 demethylation on its promoter.

To determine the contribution of BCAT1 to TKI resistance of STAC, we depleted BCAT1 expression using RNA interference. Notably, BCAT1 knockdown did not dramatically affect cell proliferation and xenograft tumor growth (Figure S6A-B) but significantly sensitized STAC-P, but not parental PC9 cells, to GEF treatment (Figure 3L-M). Meanwhile, STAC-P shBCAT1 cells restored GEF resistance after ectopic expression of shRNA-resistant BCAT1 (Figure 3L). Similarly, BCAT1 depletion also sensitized STAC-H cells to GEF (Figure S6C). Consistently, BCAT1 depletion rendered STACs xenograft tumors more sensitive to GEF (Figure 3N-P and S6D-F). Importantly, ectopic expression of BCAT1 alone indeed rendered PC9 and HCC827 cells more resistant to GEF both *in vitro* and *in vivo* (Figure S7).

To determine whether our findings from sublethal GEF treatment-induced resistance holds general meaning, we then treated PC9 cells with lethal dose of TKI following a previously established protocol (Hata et al., 2016), which is known to trigger multiple evolutionary paths of drug resistance. Through analyses of 27 established TKI-resistant subclones, we found three of them showing BCAT1 up-regulation with simultaneous H3K9me2/3 down-regulation (Figure S8A). In addition, ChIP-data confirmed that the chromatin occupancy of G9a and SUV39H1 as well as the levels of H3K9me2/3 marks at the *BCAT1* promoter were decreased in the three TKI-resistant subclones (Figure S8B-C). More importantly, BCAT1 knockdown also rendered these drug-resistant subclones more sensitive to GEF treatment (Figure S8D-F). To examine if the BCAT1-dependent mechanism of GEF resistance could also occur in the context of ERL, we established ERL-resistant PC9 cells (PC9-Erl-R) by continuous treatment of cells with sublethal dose ERL (10nM). As expected, PC9-Erl-R also exhibited increased expression of BCAT1 and strong resistance to ERL (Figure S9A-B). Moreover, knockdown of BCAT1 also sensitized PC9-Erl-R to ERL treatment (Figure S9B).

We further evaluated if the BCAT1-dependent mechanism is also relevant in targeted therapies other than EGFR inhibition. We treated ROS1-mutant HCC78 and ALK-mutant H2228 cells with sublethal dose of crizotinib continuously and established crizotinib-resistant HCC78 and H2228 cells (HCC78-Cri-R and H2228-Cri-R, respectively). Interestingly, we found that BCAT1 expression was also increased in both HCC78-Cri-R and H2228-Cri-R as compared to their parental controls (Figure S10A). Importantly, knockdown of BCAT1 also sensitized HCC78-Cri-R and H2228-Cri-R to crizotinib (Figure S10B-C). Taken together, these results demonstrate the importance of epigenetically upregulated BCAT1 in TKI resistance triggered via sublethal or lethal drug exposure.

### BCAT1 Orchestrates STAC TKI Resistance via ROS Scavenging

To determine the downstream mechanism that mediates the effects of BCAT1 on STAC drug resistance, we employed gene expression profiling analysis and found that redox-related pathway including oxidative phosphorylation and glutathione (GSH) metabolism was significantly dysregulated in BCAT1 knockdown cells (Figure S11A-B; Table S1-3). GEF treatment could promote ROS accumulation in EGFR-mutant cancer cells (Okon et al., 2015), while no obvious ROS accumulation was observed in GEF-exposed STACs (Figure S11C-D). Similarly, significant decreased ROS accumulation was detected in STAC-P xenograft tumors compared with PC9 counterparts when treated with GEF, as indicated by reduced oxidative stress marker 8-Oxo-2’-deoxyguanosine (8-OXO) (Figure S11E-F). As expected, reduced ROS accumulation was seen in PC9-Erl-R compared with PC9 parental cells when treated with ERL (Figure S9C). Likewise, we found that HCC78-Cri-R and H2228-Cri-R also showed reduced ROS levels when compared to their parental controls in response to crizotinib treatment (Figure S10D). In contrast, BCAT1 depletion in STACs increased ROS levels following GEF treatment (Figure S11G-H). Moreover, ectopic expression of BCAT1 alone reduced ROS levels in both PC9 and HCC827 cells (Figure S11I-J). These data reveal the potential link between BCAT1 and ROS scavenging in STAC.

BCAT1 expression correlates with ROS level via generation of intermediate products that suppress oxidative stress (Zhang and Han, 2017). BCAT1-engaged BCAA metabolism mainly involves two distinct pathways with different end-products: one generates glutathione (GSH) for ROS scavenging by glutamate-cysteine ligase catalytic subunit (GCLC), the rate-limiting enzyme of GSH synthesis; and the other produces coenzyme A compounds by branched-chain keto acid dehydrogenase complex (BCKDH), which participates in TCA cycle (Figure 4A) (Lu, 2013). As expected, knockdown of GCLC, but not BCKDK or BCKDHA, reduced the intracellular GSH meanwhile increased ROS level in STACs (Figure S12A-C). Notably, despite no major changes in protein levels of these enzymes between STACs and controls (Figure S3F), knockdown of GCLC clearly sensitized STACs to GEF, whereas ectopic expression of shRNA-resistant GCLC rendered GCLC-depleted cells regained GEF resistance, respectively (Figure 4B and S12D). Neither BCKDK nor BCKDHA knockdown had notable impact upon STAC TKI resistance (Figure S12E-H). More importantly, ectopic expression of GCLC completely blocked the effects of shBCAT1 on GEF resistance (Figure 4C), suggesting the requirement of GCLC in mediating BCAT1’s action.

**Figure 4.**
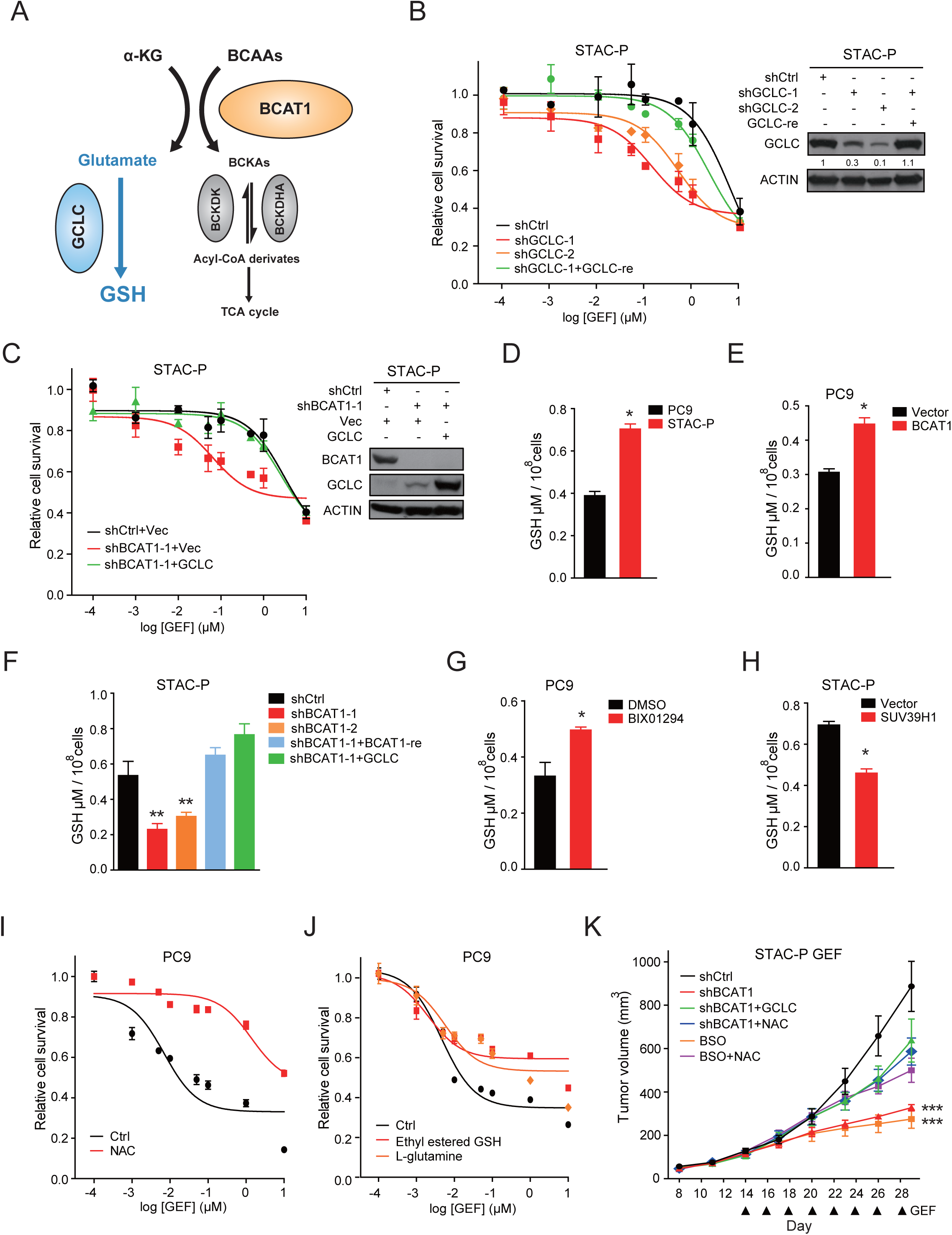
BCAA Metabolism Contributes to ROS Clearance and STAC Drug Resistance. (A) Scheme for BCAAs metabolism. (GCLC, glutamate-cysteine ligase catalytic subunit; BCKDK, branched-chain keto acid dehydrogenase complex; BCKDHA, branched chain keto acid dehydrogenase E1, alpha polypeptide; BCKA, branched-chain alpha-keto acid; α-KG, α-ketoglutarate.) (B) MTT assay of STAC-P cells with GCLC knockdown or in combination with ectopic expression of shRNA-resistant GCLC treated with indicated doses of GEF for 72 hrs. (C) MTT assay of STAC-P cells with BCAT1 knockdown or in combination with ectopic expression of Vector (Vec) or GCLC treated with indicated doses of GEF for 72 hrs. (D) GSH content of PC9 and STAC-P cells. (E) GSH content in PC9 cells with or without ectopic BCAT1 expression. (F) GSH content of STAC-P cells with BCAT1 knockdown or in combination with ectopic expression of shRNA-resistant BCAT1 or GCLC. (G) GSH content in PC9 cells treated with DMSO or BIX01294 (1µM) for 24 hrs. (H) GSH content in STAC-P cells with or without ectopic SUV39H1 expression. (I-J) MTT assay of PC9 cells treated with or without NAC (5mM) (I) or Ethyl estered GSH (100μM) or L-glutamine (10mM) (J) together with indicated doses of GEF for 72 hrs. (K) Growth curves of xenograft tumors derived from STAC-P cells with BCAT1 knockdown or in combination with ectopic expression of GCLC or dietary supplementation of NAC (40mM in drinking water) or BSO (450 mg/kg, intraperitoneal injection) following GEF treatment (50mg/kg) every other day via intraperitoneal injection. (n=6 per group). Data were presented as mean ± SEM. \**p*<0.05, \*\**p*<0.01, \*\*\**p*<0.001.

Since GCLC is involved in the biosynthesis of GSH, which serves as an important antioxidant to prevent oxidative damage caused by ROS, we then determined the role of GSH in BCAT1-mediated TKI resistance. Indeed, STACs cells displayed higher levels of GSH than their parental cells (Figure 4D and S12I). In addition, ectopic expression of BCAT1 alone increased intracellular GSH contents in both in PC9 and HCC827 cells (Figure 4E and S12J); and knockdown of BCAT1 in STACs markedly decreased intracellular GSH content (Figure 4F and S12K). Moreover, ectopic expression of shRNA-resistant BCAT1 or GCLC restored the GSH level in BCAT1-depleted STACs (Figure 4F and S12K). The contribution of BCAT1 expression to GSH synthesis was further confirmed by isotope tracing experiments (Figure S13). Consistently, BIX01294 treatment increased GSH content in PC9 and HCC827 cells (Figure 4G and S12L), and ectopic expression of SUV39H1 in STACs reduced intracellular GSH contents (Figure 4H and S12M). Since the ratio of GSH to GSSG is more indicative of the redox state of cells, we also measured the GSH/GSSG ratio in all these conditions and obtained similar results (Figure S14A-J). Further, treatment with ROS scavengers including N-acetyl cysteine (NAC), esterified GSH or L-glutamine as GSH supplement rendered PC9 cells more resistant to GEF treatment (Figure 4I-J and S14K-N). More importantly, both ectopic expression of GCLC and supplementation of NAC indeed rescued the shBCAT1 phenotype in STAC-P xenografts following GEF treatment (Figure 4K), suggesting that the sensitization of BCAT1 knockdown tumors to GEF is ROS-dependent. Collectively, these findings support the notion that BCAT1 contributes to STAC TKI resistance potentially through attenuation of ROS accumulation via GSH synthesis.

### Combination of GEF with ROS-Inducing Agents Overcomes STAC TKI Resistance

To assess the translational significance, we treated STACs combing GEF with ROS-inducing agents. Small molecular inhibitors including Piperlongumine (PL) and Phenethyl Isothiocyanate (PEITC) are known to exert anti-tumor activities through promoting ROS accumulation (Liu et al., 2014; Xiao et al., 2010). Notably, combination of GEF with PL or PEITC effectively overcame STAC-P TKI resistance *in vitro* (Figure 5A-B and S15A). Consistently, only GEF and PL combinational treatment significantly inhibited STAC-P xenograft growth and cell proliferation with excessive ROS accumulation (Figure 5C-F). Similar results were also observed in STAC-H cells (Figure S15B-C).

**Figure 5.**
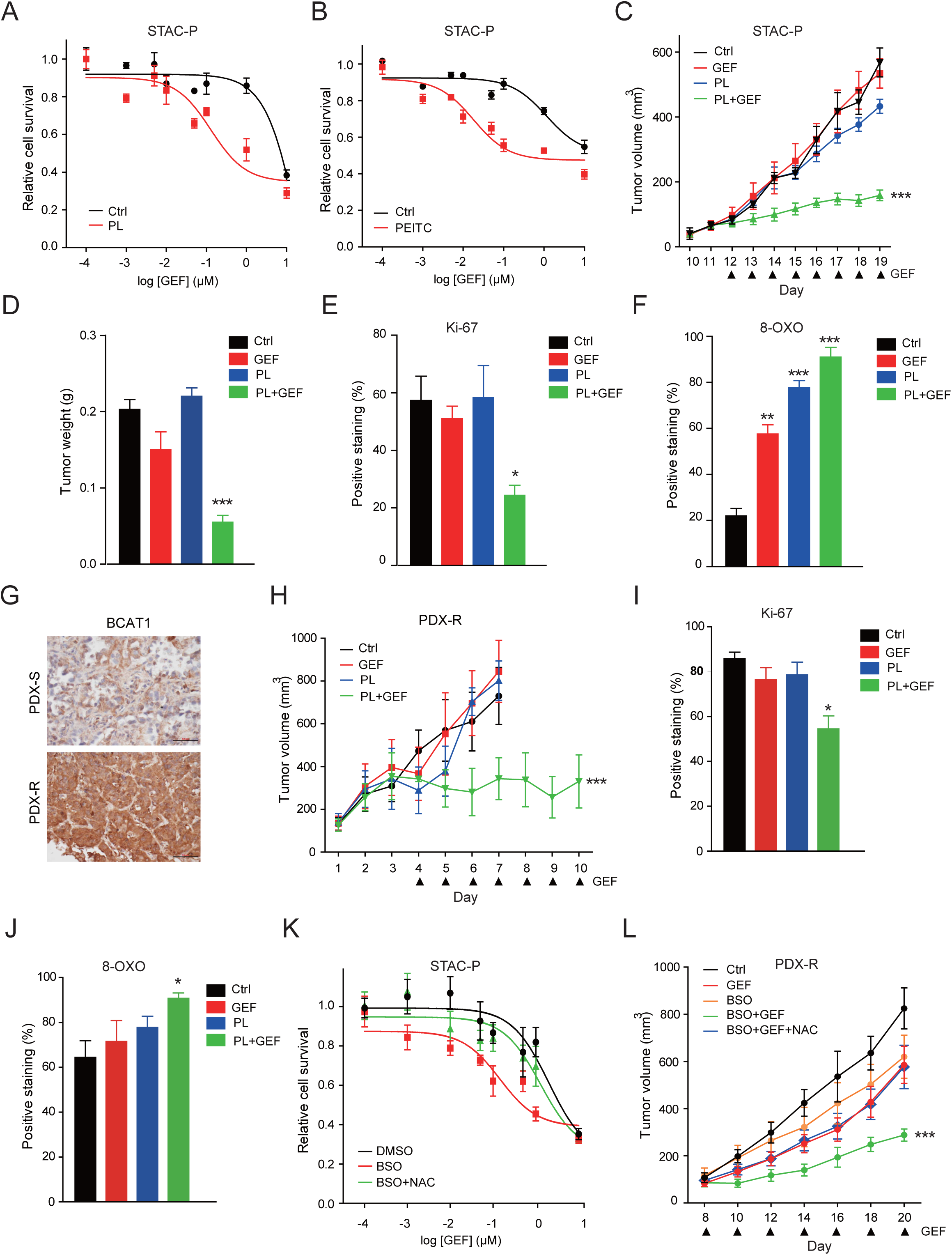
Combined GEF and PL Treatment Overcomes STAC Drug Resistance. (A-B) MTT assay of STAC-P cells treated with or without PL (0.1µM) (A) or PEITC (1µM) (B) together with indicated doses of GEF for 72 hrs. (C) Growth curves of STAC-P xenograft tumors treated with GEF (3.25mg/kg), PL (3mg/kg), or both. (n=6 per group) (D) Tumor weight for STAC-P xenografts treated as in Figure 5C. (E-F) Percentage of Ki-67 (E) and 8-OXO (F) positive staining cells in STAC-P xenografts treated as in Figure 5C. (G) Representative IHC staining of BCAT1 in EGFR TKI sensitive PDX tumor (PDX-S) and TKI-resistant PDX tumor (PDX-R). Scale bar, 50μm. (H) Growth curves of PDX-R tumors treated with GEF (50mg/kg), PL (3mg/kg), or both. (n=6 per group) (I-J) Percentage of Ki-67 (I) and 8-OXO (J) positive staining cells in PDX-R tumors treated as in Figure 5H. (K) MTT assay of STAC-P cells treated with or without BSO (200µM) or NAC (5mM) together with indicated doses of GEF for 72 hrs. (L) Growth curves of PDX-R tumors treated with GEF (50mg/kg), BSO (450 mg/kg, intraperitoneal injection) or dietary supplementation of NAC (40mM in drinking water). (n=6 per group) Data were presented as mean ± SEM. \**p*<0.05, \*\**p*<0.01, \*\*\**p*<0.001.

In addition, we established a patient-derived xenograft (PDX-resistant, PDX-R) from a patient who contained EGFR L858R mutation but showed EGFR TKI resistance. Interestingly, this PDX-R tumor showed higher BCAT1 expression compared with PDX-S (PDX-sensitive) tumor, which was sensitive to GEF treatment (Figure 5G). Similarly, only PL/GEF combinational treatment efficiently suppressed the PDX-R tumor growth, inhibited cell proliferation and promoted ROS accumulation (Figure 5H-J). To further verify our findings, we pharmacologically inhibited GSH synthesis using a more specific inhibitor, buthionine sulfoximine (BSO), which has been widely used *in vivo* with no observed toxicity and has a more specific mechanism of action. Indeed, BSO treatment sensitized STACs as well as PDX-R to GEF both *in vitro* and *in vivo*, which effects could be abolished by NAC supplementation (Figure 4K, 5K-L and S15D). Thus, these data suggest that combining TKI with ROS-inducing agents may be effective to overcome TKI resistance.

### Clinical Correlation of BCAT1 Expression with Therapeutic Response to EGFR TKI

To evaluate the association between BCAT1 expression and EGFR TKI therapeutic response, we examined 119 human lung cancer specimens with TKI-sensitive EGFR mutations including 80 biopsy samples before TKI therapy (the baseline) and 39 TKI treatment relapsed samples (Table S4). Immunohistochemistry analysis revealed a negative correlation between BCAT1 and H3K9me2 or 8-OXO level (Figure 6A-C; Table S4). Importantly, BCAT1 expression were higher in the relapsed group than the baseline group (Figure 6D), while no major difference in *BCAT1* gene copy number was observed (Figure S16). In the baseline group, high BCAT1 expression was associated with unfavorable therapeutic response to TKI treatment (Figure 6E and Table S5). Patients in the baseline group who had high-BCAT1 tumors had a shorter progression-free survival than those who had low-BCAT1 tumors (Figure 6F and Table S6). While patients in the relapsed group who did not harbor known genetic alterations, e.g., T790M mutation, MET or HER2 amplification, tended to have higher BCAT1 expression (Figure 6G). Together, these results support the correlation of BCAT1 expression with TKI response of patients with EGFR-mutant lung cancer.

**Figure 6.**
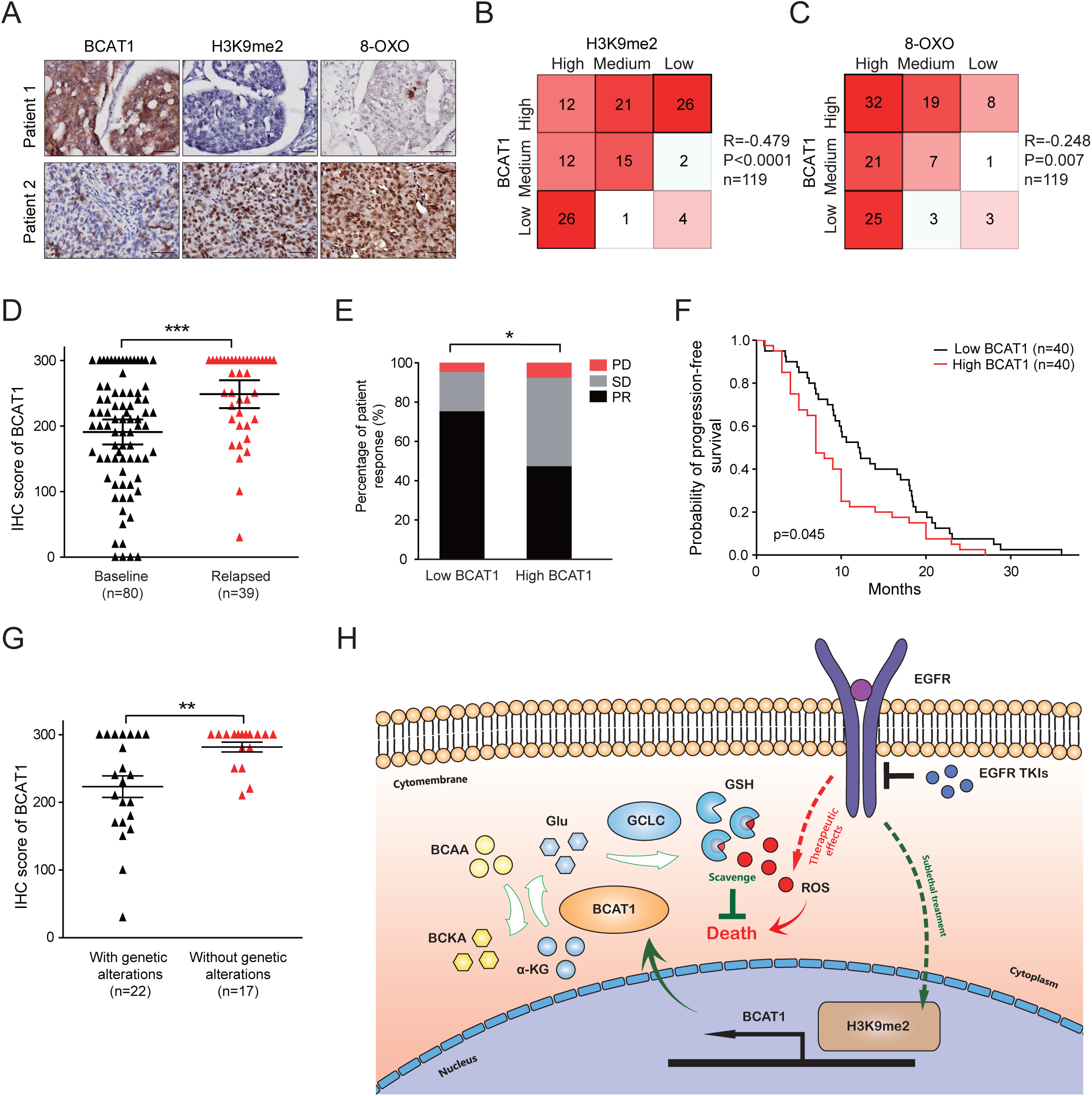
Clinical Correlation Between BCAT1 Expression and EGFR TKI Resistance. (A) Representative IHC staining for BCAT1, H3K9me2, and 8-OXO in two human EGFR-mutant lung cancer specimens. Scale bar, 50μm. (B-C) Kendall’s tau correlation analyses of IHC staining for BCAT1 with H3K9me2 (B) or with 8-OXO (C) in human EGFR-mutant lung cancer specimens. (D) Comparison of IHC score of BCAT1 expression in the baseline group and the relapsed group. Data were presented as mean ± SEM. (E) Percentage of patent response in the baseline samples according to BCAT1 expression status. (PR, partial response; SD, stable disease; and PD, progressive disease) (F) Kaplan-Meier curves for progression-free survival of patients in the baseline group according to BCAT1 expression status. (G) Comparison of IHC score of BCAT1 expression in the relapsed group with or without known genetic alterations. Data were presented as mean ± SEM. (H) A schematic diagram of BCAT1-mediated ROS scavenging in EGFR TKI resistance induced by continuous mild TKI treatment. \**p*<0.05, \*\**p*<0.01, \*\*\**p*<0.001.

## DISCUSSION

Here we report that sublethal TKI exposure for 2 hours can elicit a transient drug tolerant state in EGFR-mutant lung cancer cells. Continuous sublethal TKI exposure reinforces this tolerance and establishes persistent drug resistance through a mechanism involving epigenetically regulated BCAA metabolic reprogramming that eliminates detrimental oxidative stress (Figure 6H). It has been shown that epigenetic regulation such as H3K4 demethylation, contributes to EGFR TKI resistance triggered by lethal drug exposure through IGF-1R signaling. A recent work also proposes that lethal TKI exposure promotes drug resistance through chromatin repression via H3K9 and H3K27 methylation (Guler et al., 2017). We demonstrate that sublethal GEF exposure-induced drug resistance is potentially mediated through H3K9 demethylation. Of note, STACs established in our study are induced via continuous sublethal TKI treatment, whereas the drug-tolerant cells reported by others are established through lethal drug exposure (Guler et al., 2017; Sharma et al., 2010). The discrepancy of histone H3 methylation patterns in different TKI-resistant cells may be attributed to the different experimental systems employed. In clinical practice, patients are routinely given maximum tolerated dose of TKI and cancer evolves under sustained maximal therapeutic pressure. Interestingly, accumulative evidence has demonstrated that not all the tumor cells are evenly exposed to a lethal dose of drug and certain cancer cells may be exposed to sublethal doses of drug due to intratumoral heterogeneity resulting from complex tumor microenvironment that restrains drug diffusion and distribution (Fuso Nerini et al., 2014; Tredan et al., 2007). In fact, we did observe a heterogeneity of drug distribution within a single tumor and the activation of EGFR signaling was also highly heterogeneous in different regions of the EGFR-mutant GEF-resistant lung tumor samples (Figure S17). Our data suggest that sublethal exposure to TKI may instead enable tumor cells to develop adaptive programs to survive upon subsequent cytotoxic doses of targeted therapy.

We demonstrate that H3K9 demethylation contributes to STAC TKI resistance potentially through epigenetic upregulation of the BCAA catabolic enzyme BCAT1. Aberrant BCAT1 expression and BCAA metabolism has recently been documented to promote proliferation and progression of multiple malignancies (Hattori et al., 2017; Mayers et al., 2016; Tonjes et al., 2013). Although we cannot rule out the potential involvement of other genes in TKI resistance, our gain-of-function and loss-of-function experiments consistently support the notion that increased BCAT1 expression may be one of the important mechanisms underlying TKI resistance. Though our results were obtained from sublethal TKI-exposure model, we found that about 10% (3/27) of the TKI-resistant PC9 subclones established through lethal dose exposure also show reduced G9a and SUV39H1 binding and H3K9me2/3 marks at the *BCAT1* promoter with simultaneous BCAT1 upregulation and are indeed vulnerable to BCAT1 knockdown. These findings indicate that even during the lethal TKI treatment, a small fraction of tumor cells may evolve to become resistant via the mechanism similar to STACs, and that targeting BCAT1 may represent a potential therapeutic strategy to abrogate BCAT1-dependent drug resistance. The observations that crizotinib-resistant cells also harbored increased BCAT1 expression and that BCAT1 depletion rendered them more sensitive to crizotinib indicate that the BCAT1-dependent mechanism that we described here might be, at least in part, relevant in several other TKI resistance settings in addition to EGFR TKI resistance.

Our data suggest the importance of BCAT1-mediated metabolic reprogramming for generating GSH in mediating STAC drug resistance, which is in accordance with previously reported effects of GSH and GCLC on therapeutic resistance (Benhar et al., 2016; Zheng et al., 2016). Importantly, combining ROS-inducing agents with TKI is effective to suppress tumor growth in both STACs xenograft and PDX models, raising the possibility that adjunctive therapies designed to induce ROS accumulation might provide a potential approach to improve EGFR TKI response. Of note, non-selective ROS-inducing agents that globally increase oxidants in cancer have limited therapeutic index due to excessive toxicity in sensitive tissues such as the liver and the nervous system, and many therapeutic approaches targeting intracellular ROS levels have yielded mixed results (Chio and Tuveson, 2017; Yang et al., 2018). Given the complicated role of ROS-inducing agents in cancer therapy, more efforts are needed to verify the potential translational significance of our findings and search for specific inhibitors with less toxicity, e.g., those specifically target the BCAT1-engaged pathway other than non-selective ROS-inducing agents.

We observe an inverse correlation of BCAT1 expression with levels of H3K9me2 or oxidative stress in clinical tumor samples harboring EGFR mutations. Moreover, high BCAT1 expression is associated with unfavorable therapeutic response to EGFR TKI and shortened progression-free survival of EGFR-mutant patients, which is in agreement with our *in vitro* and *in vivo* findings. Taken together, our results identify a mechanism of EGFR TKI resistance involving adaptive response potentially through epigenetically regulated BCAT1-mediated metabolic reprogramming, which may contribute to a better understanding of the complexity and heterogeneity of drug resistance in clinic.

## LIMITATIONS OF STUDY

Our data demonstrate that STAC maintains a low level of H3K9 methylation potentially through inhibition of H3K9 methyltransferase activities. How the activities of these methyltransferases are compromised in STACs requires future efforts to dissect into details. In addition, considering that acquired drug resistance may involve diverse molecular mechanisms that arise within same patient, it is currently impractical to define which human EGFR-mutant tumor develops TKI resistance through the mechanism reported here.

## STAR★METHODS

Detailed methods are provided include the following:

- **KEY RESOURCES TABLE**
- **CONTACT FOR REAGENT AND RESOURCE SHARING**
- **EXPERIMENTAL MODEL AND SUBJECT DETAILS**
  - Cell Culture Studies
  - Animal Studies
  - Human Samples
- **METHOD DETAILS**
  - Cell Proliferation Assay
  - Cell Survival Assay
  - TKI Pre-treatment Experiments
  - Generation of Sublethal TKI Adapted Cells (STAC)
  - Plasmid Construction and Virus Infection
  - Gene Amplification Analyses
  - Real-time PCR Analyses
  - Western Blot Assay
  - Immunohistochemical Staining
  - H3K9 Methyltransferase Enzyme Activity Test
  - Chromatin Immunoprecipitation Assay
  - GSH Density Measurement
  - Intracellular Reactive Oxygen Species (ROS) Detection
  - UHPLC-qTOF-MS Analysis
  - Quantification of [13C5]-Glutamate in Condition Medium
  - Bioinformatics Analysis
  - PDX Model Establishment
  - Clinical Character Definition
- **QUANTIFICATION AND STATISTICAL ANALYSIS**
- **DATA AND SOFTWARE AVAILABILITY**

## SUPPLEMENTAL INFORMATION

Supplemental Information includes 17 figures and 6 tables and can be found in the Supplemental files.

## ACKNOWLEDGEMENTS

We are grateful for helpful comments and materials supports from Drs. Jeffrey A. Engelman, Degui Chen, Wenyi Wei and Qing Yan. This work was supported by the National Basic Research Program of China (grants 2017YFA0505501 to H.J.); the Strategic Priority Research Program of the Chinese Academy of Sciences (grants XDB19020000 to H.J.); the National Natural Science Foundation of China (grants 81430066 to H.J., 91731314 to H.J.,31621003 to H.J.,81872312 to H.J.,81871875 to L.H.,81802279 to H.H.); the Science and Technology Commission of Shanghai Municipality (grant 15XD1504000 to H.J.); and the China Postdoctoral Science Foundation (2016M601667 to H.H.).

## AUTHOR CONTRIBUTIONS

H.J. and Y.W. conceived the project. H.J., Y.W., J.Z., L.H. and D.S. designed experiments. Y.W., J.Z. and D.S. carried out most of the experiments and analyzed the data. S.R. and C.Z. provided and analyzed clinical samples. T.J. helped in ARMS analysis. Z.J., L.D., C.L. and L.C. helped in bioinformatics analysis. H.H., F.L., H.W., C.Z., K.W., L.Y., Y.J., X.H., S.H., C.G., F.L., D.G., J.Q., and D.M.G. provided technical supports and comments. H.J., L.H., and Y.W. wrote the manuscript.

