## Supplementary material for "Branched-Chain Amino Acid Metabolic Reprogramming Orchestrates Drug Resistance to EGFR Tyrosine Kinase Inhibitors": Suppl figures and methods

### SUPPLEMENTARY FIGURE LEGENDS

**Figure S1. Sublethal EGFR TKI treatment induces short-term tolerance and long-term resistance in human EGFR-mutant lung cancer cells.**

(A) PC9 cells were treated with indicated doses of GEF for 24 hrs followed by FACS analysis of PI/Annexin V positive staining.

(B) MTT assay of SH450 cells treated as in Figure 1D.

(C) PC9 and HCC827 cells were exposed to GEF (0.1μM or 1μM) for 2 hrs followed by re-exposure to indicated doses of GEF for 0.5 hr and then cultured in drug-free medium for additional 72 hr before subjected to MTT assay.

(D) PC9 cells were exposed to sublethal erlotinib (ERL) (10nM) for 2 hrs followed by recovery in drug-free medium with different times before re-exposed to indicated doses of GEF for 0.5 hr and then cultured in drug-free medium for additional 72 hrs before subjected to MTT assay.

(E) Statistical analysis of $IC_{50}$ value of PC9 and STAC-P cells (left) or HCC827 and STAC-H cells (right) in response to GEF treatment.

(F) Representative IHC staining of pEGFR (Y1068) in PC9 and STAC-P xenograft tumors treated as in Figure 1I. Scale bar, 50μm. Statistical analyses of pEGFR level were presented as mean ± SEM.

(G) Growth curves of xenograft tumors derived from HCC827 and STAC-H cells treated with or without GEF (50mg/kg) daily via intraperitoneal injection. (n=6 per group)

(H) Tumor weight for xenografts treated as in Figure S1G. Data were presented as mean ± SEM.*$p < 0.05$, **$p < 0.01$, ***$p < 0.001$.

**Figure S2. Studies of known TKI resistance mechanisms in STAC.**

(A) Detection of EGFR and KRAS mutation in PC9, STAC-P, HCC827 and STAC-H cells by ARMS method.

(B) Relative growth of PC9 and STAC-P cells were determined by MTT assay.

(C) Cell cycle analysis of PC9 and STAC-P cells. Data were presented as mean ± SEM.

(D) Representative results for phospho-RTK array. Dots on top right corner were served as internal controls.

(E) Quantitative real-time PCR analysis of indicated gene expression after shRNA knockdown in STAC-P cells.

(F) MTT assay of STAC-P cells with or without indicated gene knockdown treated with indicated doses of GEF for 72 hrs. ** $p < 0.01$, *** $p < 0.001$.

**Figure S3. The involvement of H3K9 demethylation in STAC TKI resistance.**

(A) Cells treated as in Figure 2A were harvested and subjected to western blot assay and probed with indicated antibodies.

(B) Histone demethylases (HDMT) activities for H3K9 in indicated cells. Data were presented as mean ± SEM.

(C) Relative mRNA levels of indicated H3K9 demethylases in PC9 and STAC-P cells were determined by qPCR method.

(D) Knockdown efficiency of indicated H3K9 demethylases in STAC-P cells was determined by qPCR method.

(E) Growth inhibition rate of STAC-P cells with indicated gene knockdown was determined by MTT assay following GEF treatment (10nM) for 72 hrs.

(F) Protein levels of indicated genes were determined by Western blot assay.

(G) Relative growth of indicated cell lines were determined by MTT assay.

(H-I) MTT assay of HCC827 cells with G9a knockdown or in combination with ectopic expression of shRNA-resistant G9a (H) or with SUV39H1 knockdown or in combination with ectopic expression of shRNA-resistant SUV39H1 (I) treated with indicated doses of GEF for 72 hrs.

(J) MTT assay of HCC827 cells treated with DMSO or BIX01294 (1μM) together with indicated doses of GEF for 72 hrs.

(K) MTT assay of STAC-H cells with or without ectopic expression of G9a or SUV39H1 treated with indicated doses of GEF for 72 hrs. $^{**}p < 0.01$, $^{***}p < 0.001$.

**Figure S4. TET may not be involved in mediating GEF resistance.**

(A-B) Dot blots detecting the 5mC (A) and 5hmC (B) using the genomic DNA extracted from the indicated cells.

(C-F) MTT assay of PC9 (C), STAC-P (D), HCC827 (E) and STAC-H (F) cells with shRNA knockdown of TET1, TET2 or TET3.

(G) Knockdown efficacy of indicated genes was determined by qPCR. $^{*}p < 0.05$.

**Figure S5. BCAT1 is epigenetically upregulated in STAC by H3K9 demethylation.**

(A) List of 22 consistently upregulated genes from the integrative analyses of microarray gene expression and RNA-seq data.

(B) Knockdown efficiency of indicated gene expression in STAC-P cells was determined by qPCR.

(C-D) Relative mRNA (C) and protein (D) levels of BCAT1 in indicated cells.

(E) Representative IHC staining of BCAT1 in PC9 and STAC-P xenograft tumors. Scale bar, 50μm. Statistical analyses of BCAT1 staining were presented as mean ± SEM.

(F) Protein levels of BCAT1 and H3K9me2 levels in PC9 cells with indicated passages continuously treated with sublethal GEF (10nM).

(G) ChIP assay of the H3K9me2 or H3K9me3 enrichment on *BCAT1* promoter relative to immunoglobulin G (IgG) in indicated cells.

(H) ChIP assay of the G9a or SUV39H1 enrichment on *BCAT1* promoter relative to IgG in indicated cells.

(I) ChIP assay of the H3K9me2 or H3K9me3 enrichment on *BCAT1* promoter relative to IgG in PC9 cells treated with DMSO or BIX01294 (1μM) for 24 hrs.

(J) Protein levels of BCAT1 in HCC827 cells with or without G9a or SUV39H1 knockdown or in combination with ectopic expression of shRNA-resistant cDNAs.

(K) Protein levels of BCAT1 in STAC-H cells with or without ectopic expression of G9a or SUV39H1.

(L) Protein levels of BCAT1 in HCC827 cells treated with DMSO or BIX01294 (1μM) for 24 hrs. *$p < 0.05$, **$p < 0.01$, ***$p < 0.001$.

**Figure S6. BCAT1 knockdown sensitizes STAC to GEF treatment.**

(A) Relative growth of indicated cell lines were determined by MTT assay.

(B) Growth curves of xenograft tumors derived from STAC-P cells with or without BCAT1 knockdown. (n=6 per group)

(C) MTT assay of STAC-H cells with BCAT1 knockdown alone or in combination with ectopic expression of shRNA-resistant BCAT1 treated with indicated doses of GEF for 72 hrs.

(D) Growth curves of xenograft tumors derived from STAC-H cells with BCAT1 knockdown or in combination with ectopic expression of shRNA-resistant BCAT1 following GEF treatment (50mg/kg) daily via intraperitoneal injection. (n=6 per group)

(E) Tumor weight for STAC-H xenografts treated as in Figure S6D. Plots were presented as mean ± SEM.

(F) Representative immunohistochemical (IHC) staining of Ki-67 in STAC-H xenografts treated as in Figure S6D. Scale bar, 50μm. Statistical analyses of Ki-67 staining were presented as mean ± SEM. *$p < 0.05$, **$p < 0.01$.

**Figure S7. BCAT1 expression alone confer cells more resistant to GEF treatment.**

(A-B) MTT assay of PC9 cells (A) or (B) with or without ectopic BCAT1 expression treated with indicated doses of GEF for 72 hrs.

(C) Tumor weight for xenografts derived from PC9 cells with or without ectopic BCAT1 expression following GEF treatment (50mg/kg) daily via intraperitoneal injection. (n=6 per group).

(D-E) Representative IHC staining of Ki-67 in xenograft tumors treated as in Figure S7C. Scale bar, 50μm (D). Statistical analyses of Ki-67 staining were presented as mean ± SEM (E).

(F) Tumor weight for xenografts derived from HCC827 cells with or without ectopic BCAT1 expression following GEF treatment (50mg/kg) daily via intraperitoneal injection. (n=6 per group).

(G-H) Representative IHC staining of Ki-67 in xenograft tumors treated as in Figure S7F. Scale bar, 50μm (G). Statistical analyses of Ki-67 staining were presented as mean ± SEM (H). *$p < 0.05$, **$p < 0.01$, ***$p < 0.001$.

**Figure S8. Targeting BCAT1 sensitizes regular dose-induced drug-resistant PC9 subclones to GEF treatment.**

(A) MTT assay of parental PC9 cells or three PC9 drug-resistant (PC9-DR) subclones (PC9-DR-2, PC9-DR-4 and PC9-DR-21) established through continuous GEF exposure (300nM) treated with indicated doses of GEF for 72 hrs. Cell lysates were probed with indicated antibodies.

(B-C) ChIP assay of the enrichment of H3K9me2 or H3K9me3 (B) or the enrichment of G9a or SUV39H1 (C) on *BCAT1* promoter relative to IgG in indicated cells.

(D-F) MTT assay of the three PC9-DR subclones with BCAT1 knockdown alone or in combination with ectopic expression of shRNA-resistant BCAT1 treated with indicated doses of GEF for 72 hrs. *$p < 0.05$.

**Figure S9. PC9 cells may acquire ERL resistance through a BCAT1-dependent mechanism.**

(A) Protein levels of BCAT1 in parental PC9 or ERL-resistant PC9 cells (PC9-Erl-R) with BCAT1 knockdown alone or in combination with ectopic expression of shRNA-resistant BCAT1.

(B) MTT assay of parental PC9 or ERL-resistant PC9 cells (PC9-Erl-R) with BCAT1 knockdown alone or in combination with ectopic expression of shRNA-resistant BCAT1 treated with indicated doses of ERL for 72 hrs.

(C) FACS using DCFH-DA to detect ROS level of the parental PC9 or PC9-Erl-R with DMSO or ERL (10nM) treatment for 2 hrs. Results were presented as mean ± SEM. ***$p < 0.001$.

**Figure S10. Knockdown of BCAT1 sensitizes Cri-resistant cells to Crizotinib treatment.**

(A) Protein levels of BCAT1 in parental or crizotinib-resistant HCC78 and H2228 cells.

(B) Protein levels of BCAT1 in crizotinib-resistant HCC78 and H2228 cells with or without BCAT1 knockdown.

(C) MTT assay of parental or crizotinib-resistant HCC78 or H2228 cells with or without BCAT1 knockdown treated with indicated doses of crizotinib for 72 hrs.

(D) FACS using DCFH-DA to detect ROS level of the parental or crizotinib-resistant HCC78 or H2228 cells with DMSO or crizotinib (200nM) treatment for 2 hrs. *$p < 0.05$, ***$p < 0.001$.

**Figure S11. Knockdown of BCAT1 promotes excessive ROS accumulation in STAC.**

(A-B) Enrichment plot of significantly dysregulated KEGG pathway of oxidative phosphorylation (A) and glutathione metabolism (B) in STAC-P cells with or without BCAT1 knockdown.

(C-D) FACS analyses of DCFH-DA to detect ROS level in PC9 and STAC-P cells (C) or HCC827 and STAC-H cells (D) treated with or without GEF (10nM) for 2 hrs. Results were presented as mean ± SEM.

(E) Representative IHC staining of EGFR and 8-OXO in xenograft tumors derived from PC9 and STAC-P cells treated as in Figure 1I. Scale bar, 50μm.

(F) Results of percentage of 8-OXO positive staining cells in xenograft tumors treated as in Figure 1I were presented as mean ± SEM.

(G-H) FACS analyses of DCFH-DA to detect ROS level in STAC-P (G) or STAC-H (H) cells with or without BCAT1 knockdown following GEF treatment (10nM) for 2 hrs.

(I-J) FACS analyses of DCFH-DA to detect ROS level in PC9 (I) or HCC827 (H) cells with or without ectopic expression of BCAT1 following GEF treatment (10nM) for 2 hrs. Results were presented as mean ± SEM. *$p < 0.05$, **$p < 0.01$.

**Figure S12. BCAT1 regulates GEF resistance through promoting ROS clearance.**

(A-B) FACS analyses of DCFH-DA to detect ROS level in STAC-P (A) or STAC-H (B) cells with indicated gene knockdown following GEF treatment (10nM) for 2 hrs.

(C) GSH content in STAC-P and STAC-H cells with indicated gene knockdown.

(D) MTT assay of STAC-H cells with GCLC knockdown alone or in combination with ectopic expression of shRNA-resistant GCLC treated with indicated doses of GEF for 72 hrs.

(E-F) MTT assay of STAC-P cells with or without knockdown of BCKDK (E) or BCKDHA (F) treated with indicated doses of GEF for 72 hrs.

(G-H) MTT assay of STAC-H cells with or without knockdown of BCKDK (G) or BCKDHA (H) treated with indicated doses of GEF for 72 hrs.

(I) GSH content of HCC827 and STAC-H cells.

(J) GSH content of HCC827 with or without ectopic BCAT1 expression.

(K) GSH content of STAC-H cells with BCAT1 knockdown alone or in combination with ectopic expression of shRNA-resistant BCAT1.

(L) GSH content in HCC827 cells treated with DMSO or BIX01294 (1μM) for 24 hrs.

(M) GSH content in STAC-H cells with or without ectopic SUV39H1 expression. *$p < 0.05$, **$p < 0.01$, ***$p < 0.001$.

**Figure S13. BCAT1 contributes to glutamate flux for GSH biosynthesis.**

(A-C) Isotopologue spectral analysis of $^{13}C$-labeled glutamate [M+5] (A), pyroglutamic acid [M+5] (B) and GSH [M+5] (C) in indicated cell lines using [$^{13}C_5$]-glutamine as the tracer.

(D) LC-MS/MS showed the concentration of $^{13}C$-labeled glutamate in the conditional medium of indicated cells.

(E-F) Isotopologue spectral analysis of $^{13}C$-labeled γ-glutamylcysteine [M+4] (E) and GSH [M+4] (F) in indicated cell lines using [$^{13}C_6$-$^{15}N_2$]-cystine as the tracer. *$p < 0.05$.

**Figure 14. BCAT1 is involved in GSH biosynthesis.**

(A) Ratio of GSH/GSSG in PC9 and STAC-P cells.

(B) Ratio of GSH/GSSG in HCC827 and STAC-H cells.

(C) Ratio of GSH/GSSG in PC9 cells with or without ectopic BCAT1 expression.

(D) Ratio of GSH/GSSG in HCC827 cells with or without ectopic BCAT1 expression.

(E) Ratio of GSH/GSSG in STAC-P cells with BCAT1 knockdown alone or in combination with ectopic expression of shRNA-resistant BCAT1 or GCLC.

(F) Ratio of GSH/GSSG in STAC-H cells with BCAT1 knockdown or in combination with ectopic expression of shRNA-resistant BCAT1.

(G) Ratio of GSH/GSSG in PC9 cells treated with DMSO or BIX01294 (1μM) for 24 hrs.

(H) Ratio of GSH/GSSG in HCC827 cells treated with DMSO or BIX01294 (1μM) for 24 hrs.

(I) Ratio of GSH/GSSG in STAC-P cells with or without ectopic SUV39H1 expression.

(J) Ratio of GSH/GSSG in STAC-H cells with or without ectopic SUV39H1 expression.
(K-L) GSH content in PC9 (J) or HCC827 (K) cells treated with DMSO, or Ethyl estered GSH (100μM), L-glutamine (10mM), NAC (5mM), or DL-BSO (200μM).
(M-N) Ratio of GSH/GSSG in PC9 (L) or HCC827 (M) cells treated with DMSO, or Ethyl estered GSH (100μM), L-glutamine (10mM), NAC (5mM), or DL-BSO (200μM). *$p < 0.05$, **$p < 0.01$, ***$p < 0.001$.

**Figure S15. Combined GEF and PL treatment overcomes STAC drug resistance.**

(A) MTT assay of STAC-H cells treated with or without PL (0.1μM) or PEITC (1μM) together with indicated doses of GEF for 72 hrs.
(B) Growth curves of STAC-H xenograft tumors treated with GEF (3.25mg/kg), PL (3mg/kg), or both. (n=6 per group)
(C) Tumor weight for STAC-H xenografts treated as in Figure S15B.
(D) MTT assay of STAC-H cells treated with or without BSO (200μM) or NAC (5mM) together with indicated doses of GEF for 72 hrs. ***$p < 0.001$.

**Figure S16. Relative *BCAT1* gene copy number in relapsed tumor samples and baseline samples from patients with EGFR-mutant lung cancer.**

**Figure S17. Distribution of DOX and activation of EGFR signaling in GEF-treated EGFR-mutant lung cancer samples was heterogeneous in vivo.**

(A) Representative fluorescence photos for DOX distribution in PC9 xenograft tumors with DOX treatment (5mg/kg/day). Scale bar: 50μm.
(B-C) Representative IHC staining for Ki-67 (B) and cleaved Caspase-3 (C) in PC9 xenograft tumors with DOX treatment (5mg/kg/day). Scale bar: 50μm.

(D-F) Representative IHC staining for pEGFR (D), pERK (E), and pAKT (F) in lung tumor samples from EGFR-mutant patients with or without treated with GEF (250mg daily). Scale bar: 50 μm.

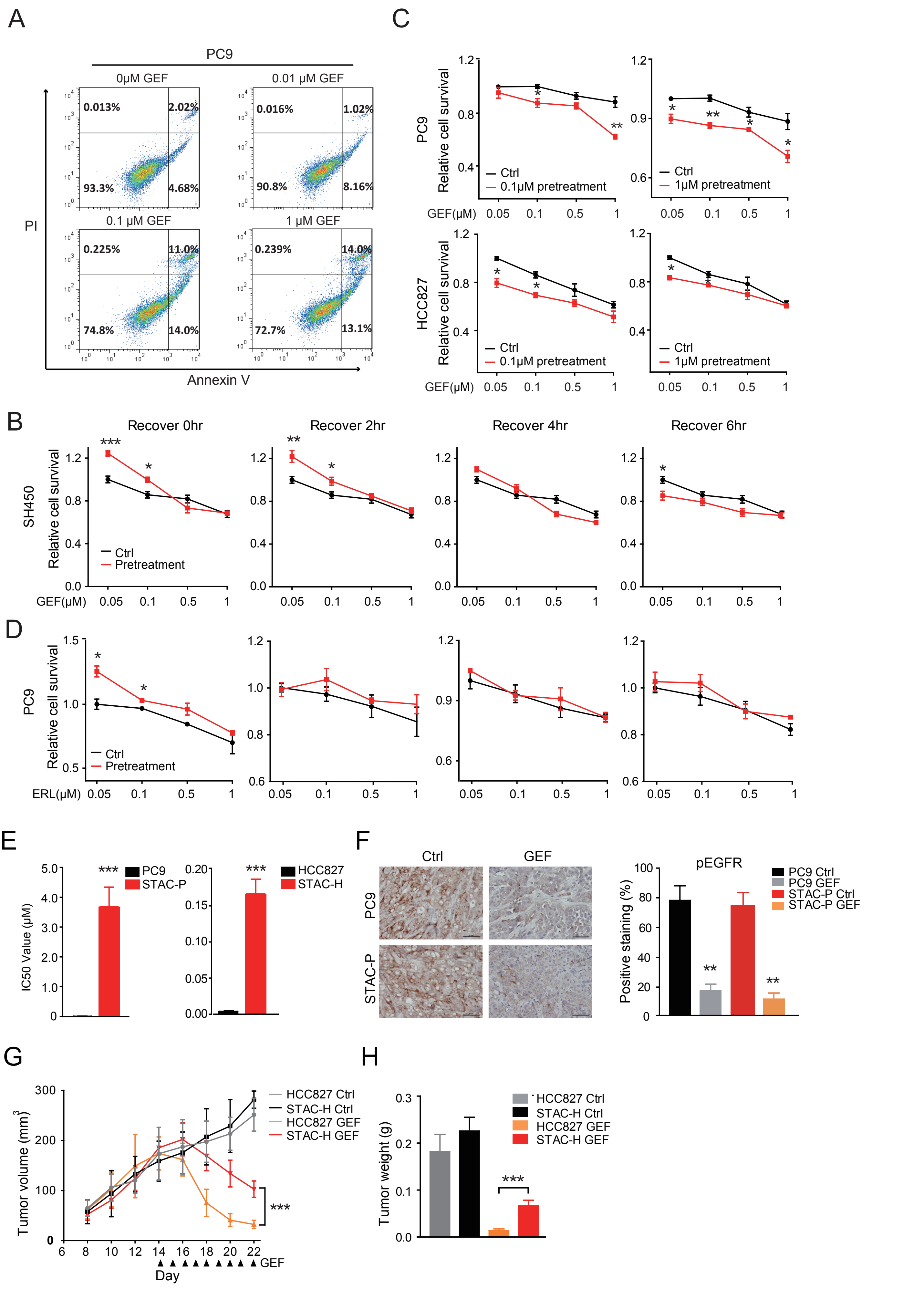

A
PC9
0μM GEF
0.01 μM GEF
0.1 μM GEF
1 μM GEF
PI
Annexin V
C
PC9
HCC827
Relative cell survival
Ctrl
0.1μM pretreatment
1μM pretreatment
GEF(μM)
B
Recover 0hr
Recover 2hr
Recover 4hr
Recover 6hr
SH450
Pretreatment
D
ERL(μM)
E
IC50 Value (μM)
PC9
STAC-P
HCC827
STAC-H
F
Ctrl
GEF
pEGFR
Positive staining (%)
PC9 Ctrl
PC9 GEF
STAC-P Ctrl
STAC-P GEF
G
Tumor volume (mm3)
HCC827 Ctrl
STAC-H Ctrl
HCC827 GEF
STAC-H GEF
Day
H
Tumor weight (g)


Figure S1

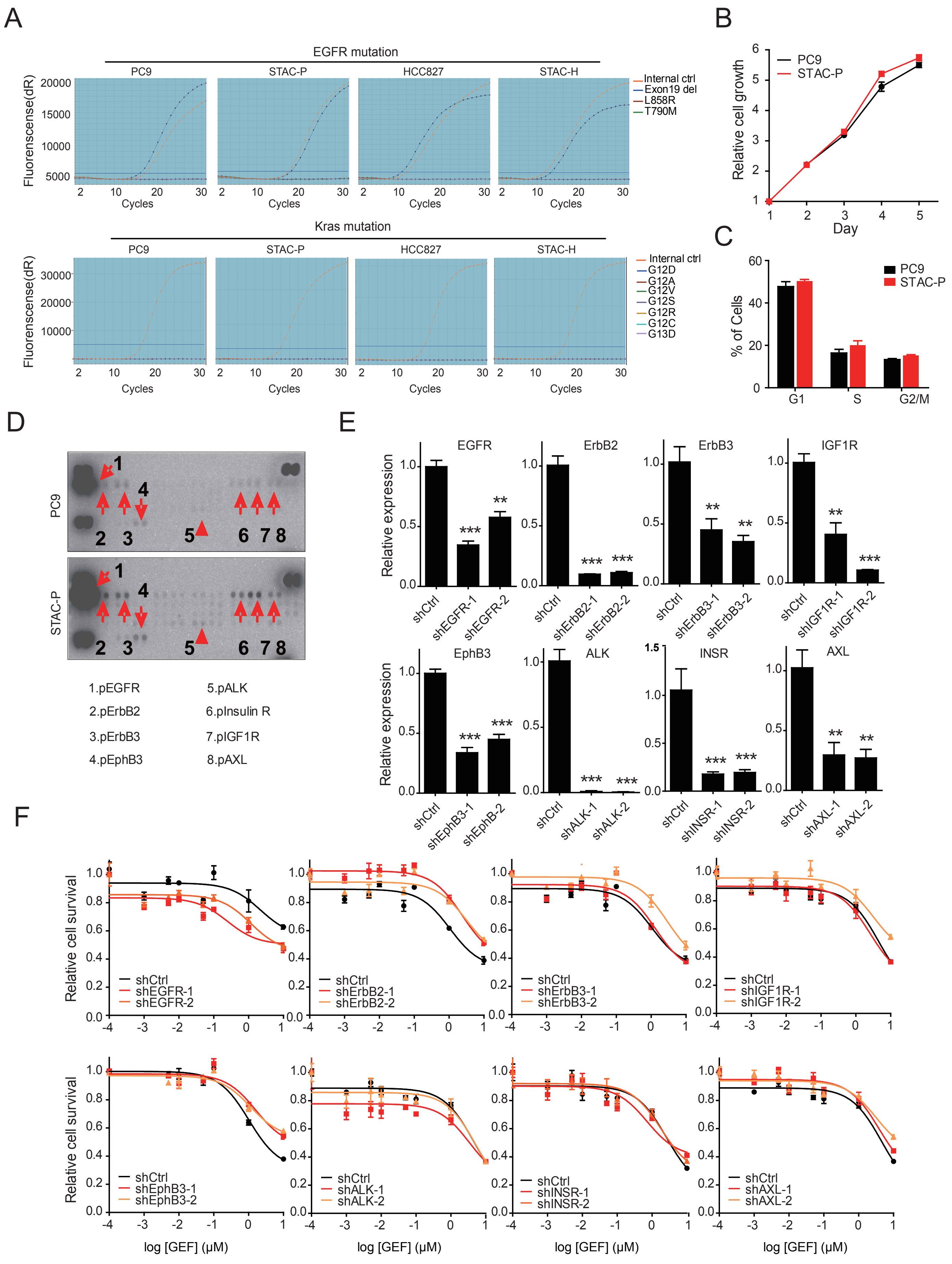
A
EGFR mutation
PC9
STAC-P
HCC827
STAC-H
Fluorenscense(dR)
Cycles
Internal ctrl
Exon19 del
L858R
T790M
Kras mutation
G12D
G12A
G12V
G12S
G12R
G12C
G13D
B
Relative cell growth
Day
C
% of Cells
G1
S
G2/M
D
1.pEGFR
2.pErbB2
3.pErbB3
4.pEphB3
5.pALK
6.pInsulin R
7.pIGF1R
8.pAXL
E
Relative expression
EGFR
ErbB2
ErbB3
IGF1R
EphB3
ALK
INSR
AXL
F
Relative cell survival
log [GEF] (μM)
shCtrl
shEGFR-1
shEGFR-2
shErbB2-1
shErbB2-2
shErbB3-1
shErbB3-2
shIGF1R-1
shIGF1R-2
shEphB3-1
shEphB3-2
shALK-1
shALK-2
shINSR-1
shINSR-2
shAXL-1
shAXL-2

Figure S2

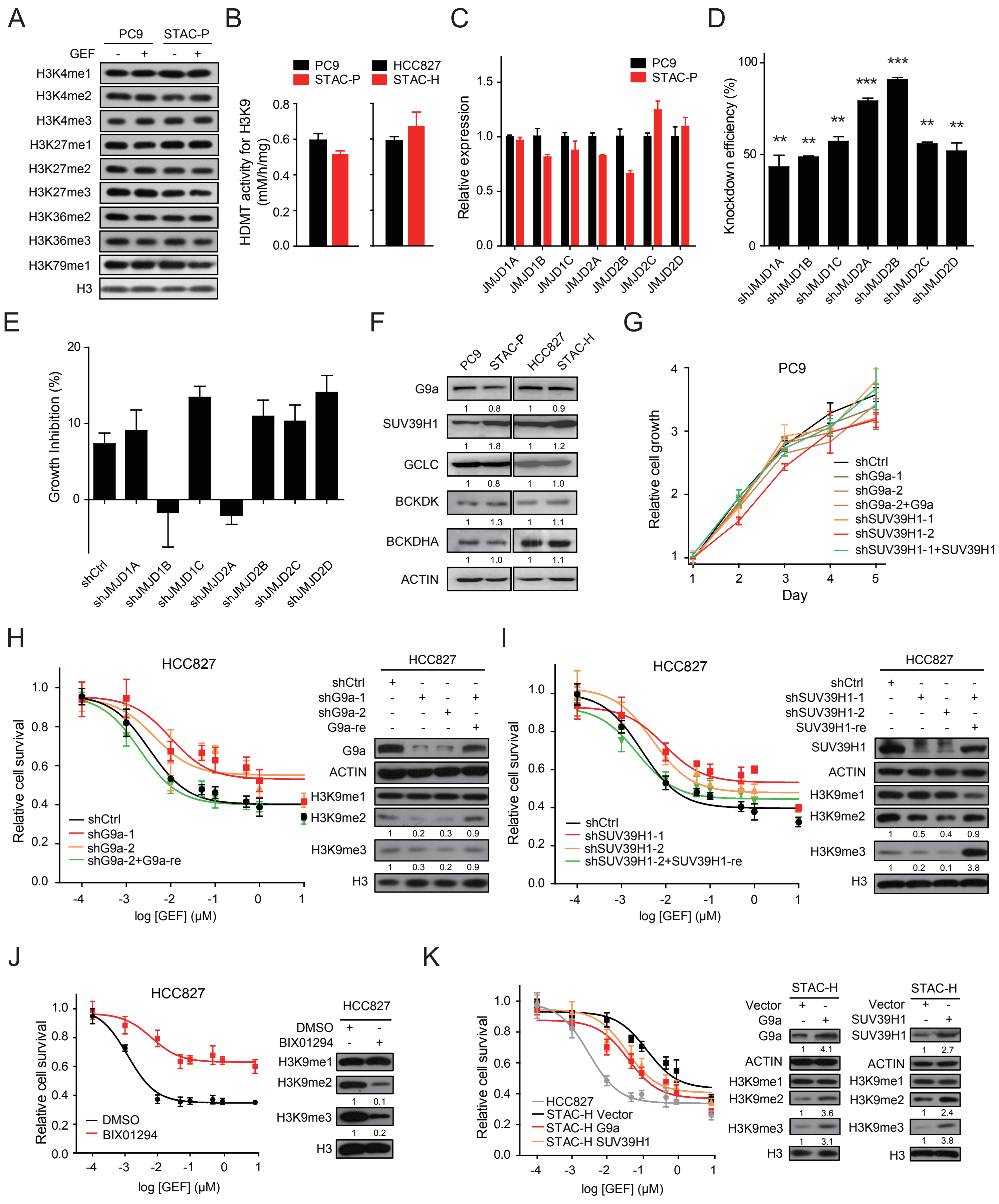
A
PC9
STAC-P
GEF
H3K4me1
H3K4me2
H3K4me3
H3K27me1
H3K27me2
H3K27me3
H3K36me2
H3K36me3
H3K79me1
H3
B
PC9
STAC-P
HCC827
STAC-H
HDMT activity for H3K9 (mM/h/mg)
C
PC9
STAC-P
Relative expression
JMJD1A
JMJD1B
JMJD1C
JMJD2A
JMJD2B
JMJD2C
JMJD2D
D
Knockdown efficiency (%)
shJMJD1A
shJMJD1B
shJMJD1C
shJMJD2A
shJMJD2B
shJMJD2C
shJMJD2D
E
Growth Inhibition (%)
shCtrl
F
PC9
STAC-P
HCC827
STAC-H
G9a
SUV39H1
GCLC
BCKDK
BCKDHA
ACTIN
G
PC9
Relative cell growth
Day
shCtrl
shG9a-1
shG9a-2
shG9a-2+G9a
shSUV39H1-1
shSUV39H1-2
shSUV39H1-1+SUV39H1
H
HCC827
Relative cell survival
log [GEF] (μM)
shG9a-2+G9a-re
G9a-re
H3K9me1
H3K9me2
H3K9me3
I
shSUV39H1-2+SUV39H1-re
SUV39H1-re
J
DMSO
BIX01294
K
HCC827
STAC-H Vector
STAC-H G9a
STAC-H SUV39H1
Vector

Figure S3

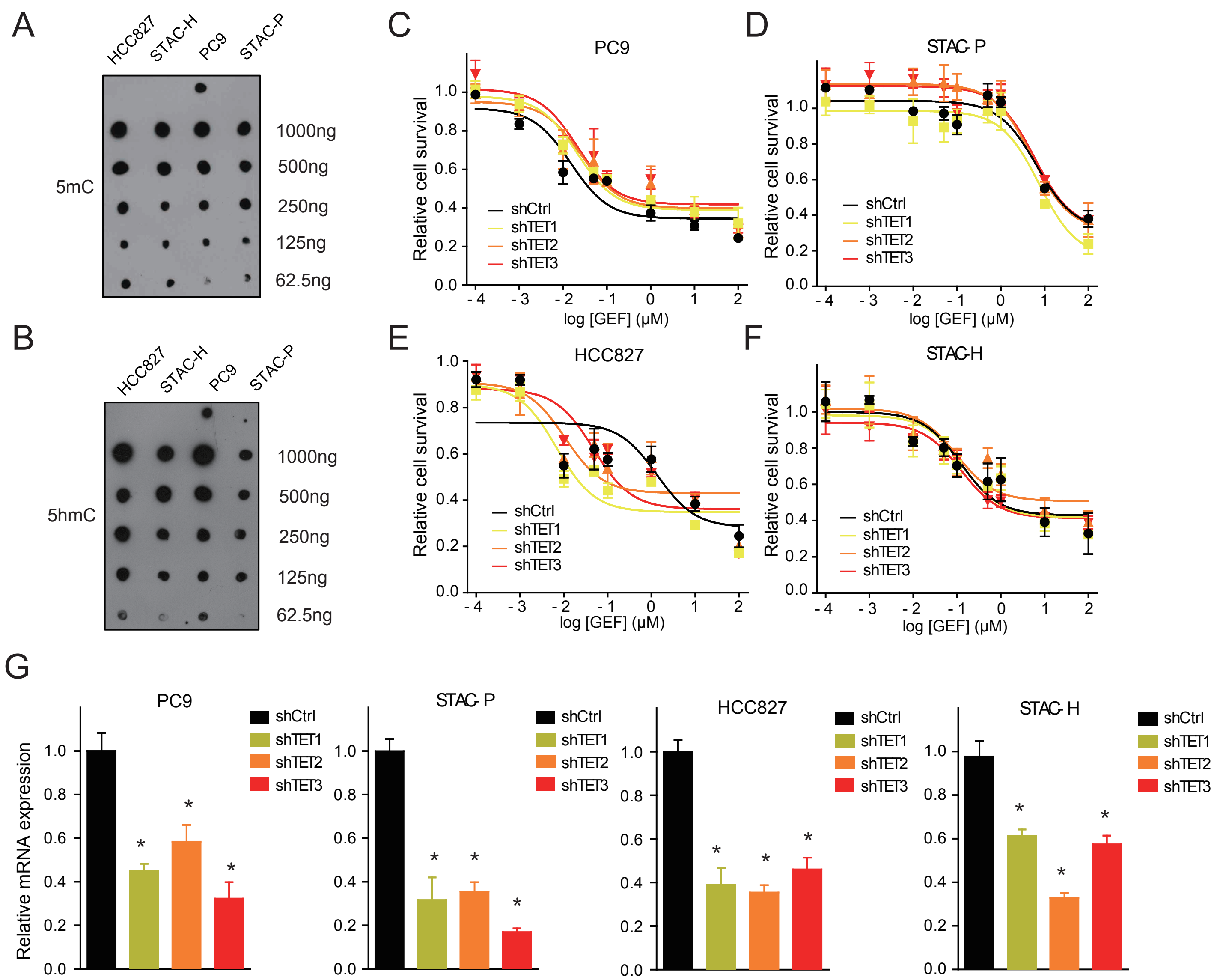

A
HCC827
STAC-H
PC9
STAC-P
5mC
1000ng
500ng
250ng
125ng
62.5ng
B
5hmC
C
PC9
D
STAC- P
E
HCC827
F
STAC-H
Relative cell survival
log [GEF] (μM)
shCtrl
shTET1
shTET2
shTET3
G
Relative mRNA expression
STAC- H


Figure S4

A

| Index | Gene_symbol | RNA_seq_logFC | Microarray_logFC |
|---|---|---|---|
| 1 | BMP5 | 5.09 | 2.44 |
| 2 | FAM171A1 | 3.74 | 1.16 |
| 3 | CYB5R2 | 3.56 | 1.73 |
| 4 | BCAT1 | 3.55 | 0.93 |
| 5 | HSD3B1 | 3.42 | 1.43 |
| 6 | RASGRF1 | 3.24 | 0.95 |
| 7 | 4-MAR | 3.07 | 1.09 |
| 8 | IL1RL1 | 2.80 | 0.90 |
| 9 | DENND2A | 2.53 | 1.85 |
| 10 | PRR5L | 2.37 | 1.12 |
| 11 | IL11 | 2.02 | 0.90 |
| 12 | FLNC | 2.01 | 1.79 |
| 13 | LAMC2 | 2.00 | 1.10 |
| 14 | HBEGF | 1.98 | 1.05 |
| 15 | CTGF | 1.97 | 1.67 |
| 16 | MSN | 1.90 | 1.33 |
| 17 | CYR61 | 1.51 | 1.31 |
| 18 | TMEM171 | 1.47 | 1.56 |
| 19 | AXL | 1.38 | 1.26 |
| 20 | NUP214 | 1.37 | 1.08 |
| 21 | EPB41L2 | 1.27 | 1.05 |
| 22 | NCF2 | 1.21 | 1.22 |

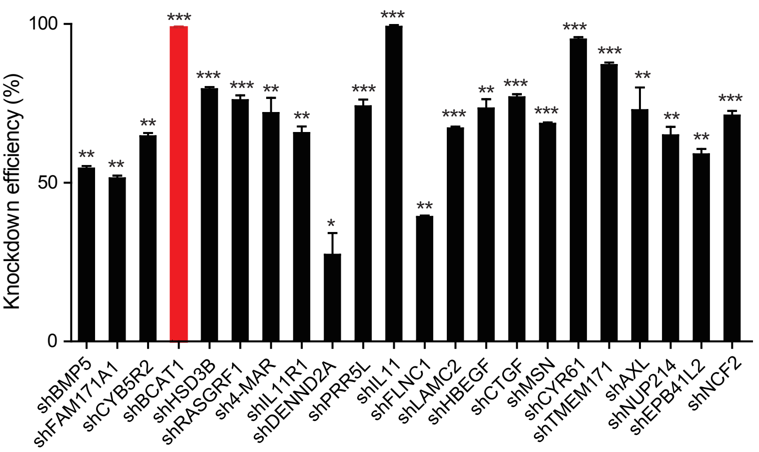


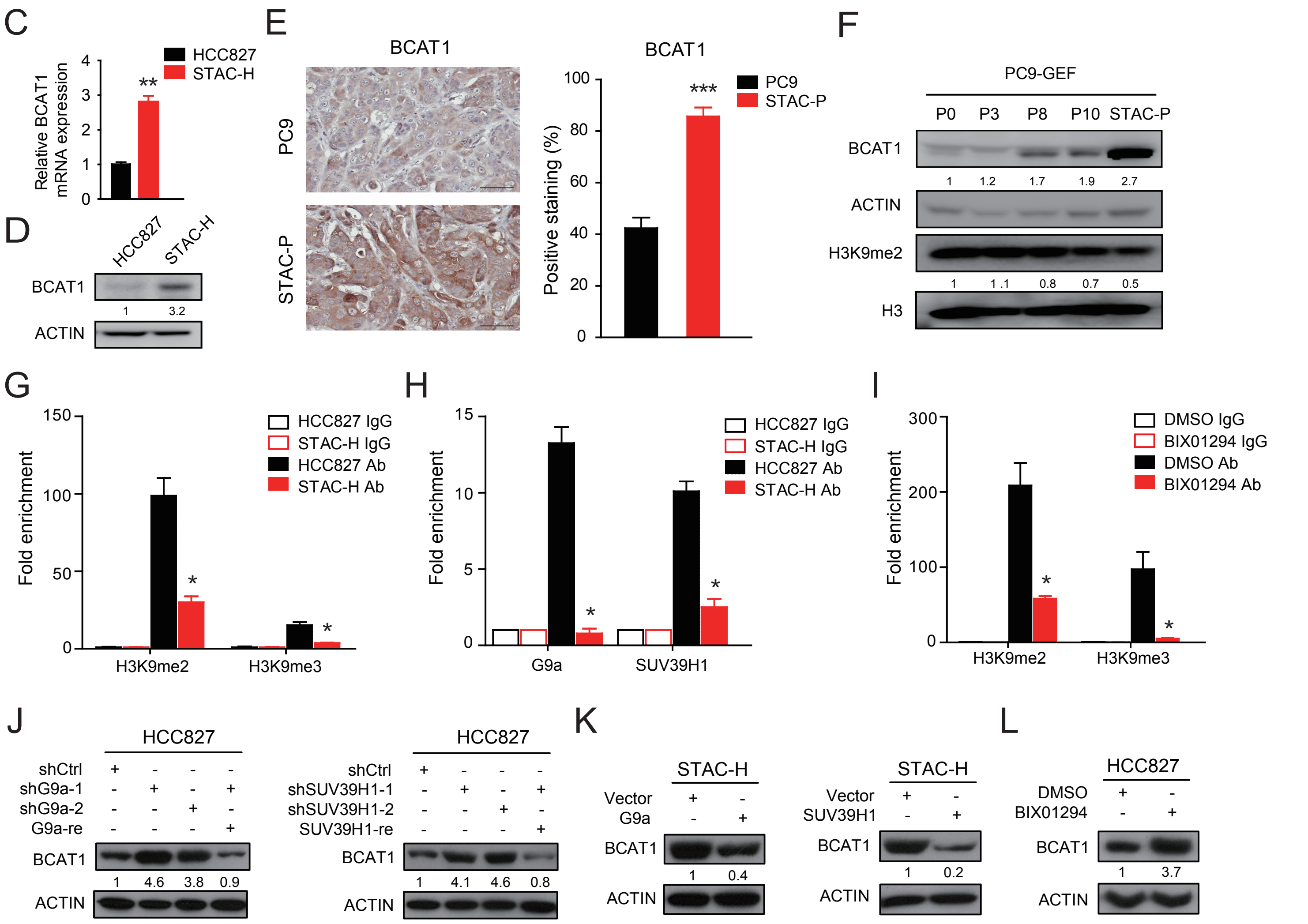


Figure S5

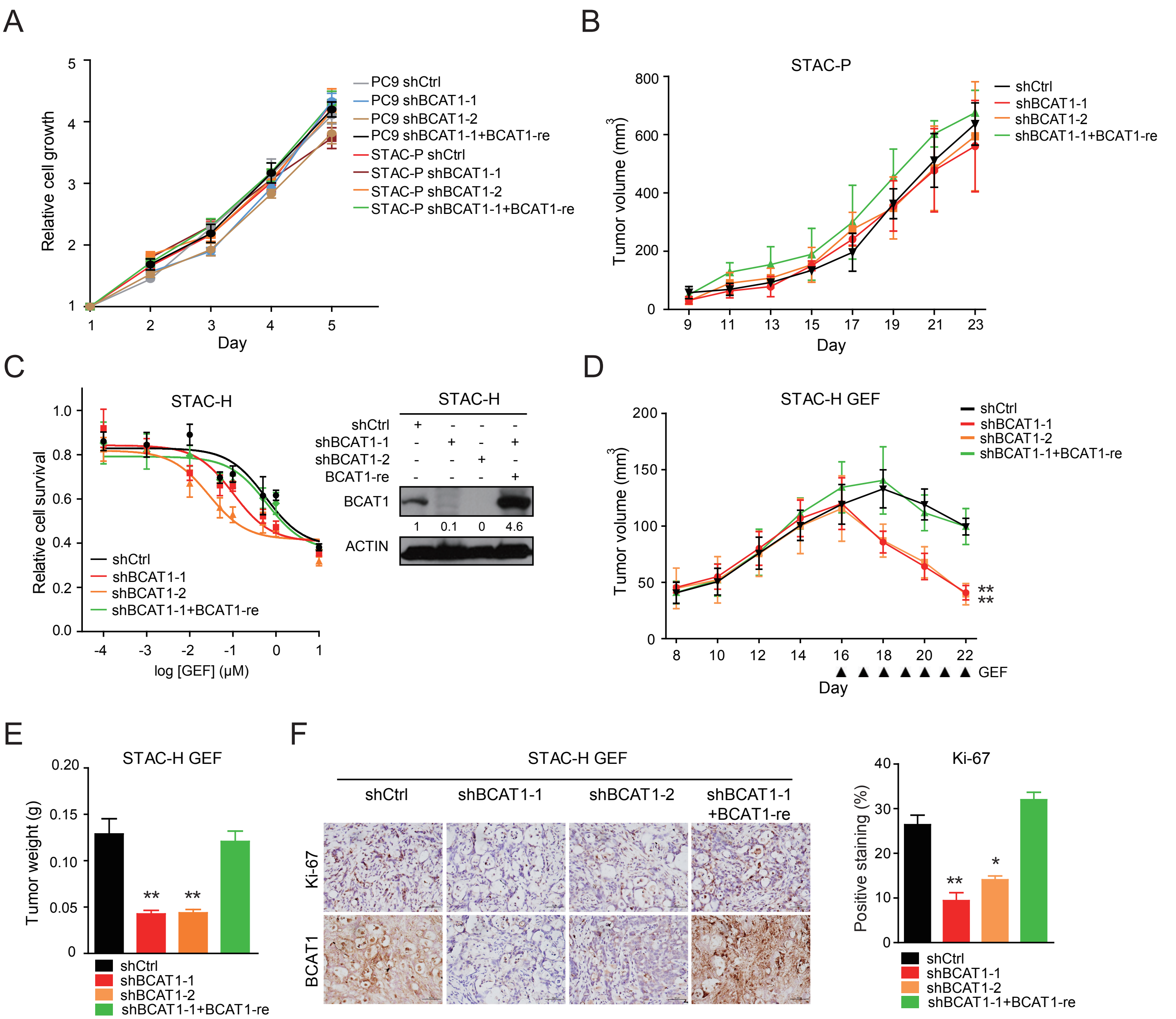
A
Relative cell growth
Day
PC9 shCtrl
PC9 shBCAT1-1
PC9 shBCAT1-2
PC9 shBCAT1-1+BCAT1-re
STAC-P shCtrl
STAC-P shBCAT1-1
STAC-P shBCAT1-2
STAC-P shBCAT1-1+BCAT1-re
B
STAC-P
Tumor volume (mm3)
Day
shCtrl
shBCAT1-1
shBCAT1-2
shBCAT1-1+BCAT1-re
C
STAC-H
Relative cell survival
log [GEF] (μM)
shCtrl
shBCAT1-1
shBCAT1-2
shBCAT1-1+BCAT1-re
STAC-H
shCtrl
shBCAT1-1
shBCAT1-2
BCAT1-re
BCAT1
1 0.1 0 4.6
ACTIN
D
STAC-H GEF
Tumor volume (mm3)
Day
GEF
shCtrl
shBCAT1-1
shBCAT1-2
shBCAT1-1+BCAT1-re
E
STAC-H GEF
Tumor weight (g)
shCtrl
shBCAT1-1
shBCAT1-2
shBCAT1-1+BCAT1-re
F
STAC-H GEF
shCtrl
shBCAT1-1
shBCAT1-2
shBCAT1-1
+BCAT1-re
Ki-67
BCAT1
Ki-67
Positive staining (%)
shCtrl
shBCAT1-1
shBCAT1-2
shBCAT1-1+BCAT1-re

Figure S6

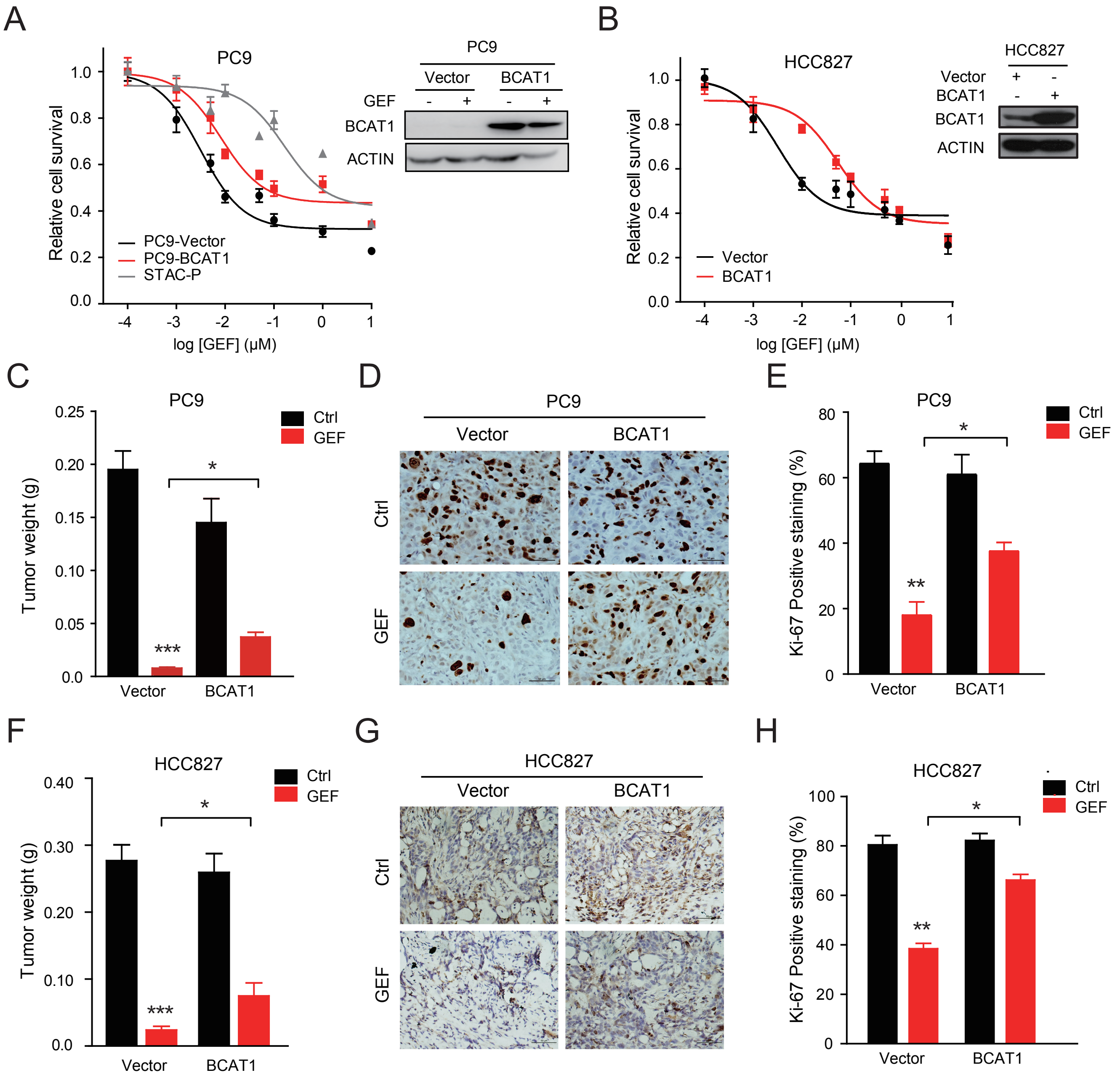
A
PC9
Relative cell survival
log [GEF] (μM)
PC9-Vector
PC9-BCAT1
STAC-P
Vector
BCAT1
GEF
BCAT1
ACTIN
B
HCC827
Relative cell survival
log [GEF] (μM)
Vector
BCAT1
ACTIN
C
PC9
Tumor weight (g)
Ctrl
GEF
Vector
BCAT1
D
PC9
Vector
BCAT1
Ctrl
GEF
E
PC9
Ki-67 Positive staining (%)
Ctrl
GEF
Vector
BCAT1
F
HCC827
Tumor weight (g)
Ctrl
GEF
Vector
BCAT1
G
HCC827
Vector
BCAT1
Ctrl
GEF
H
HCC827
Ki-67 Positive staining (%)
Ctrl
GEF
Vector
BCAT1

Figure S7

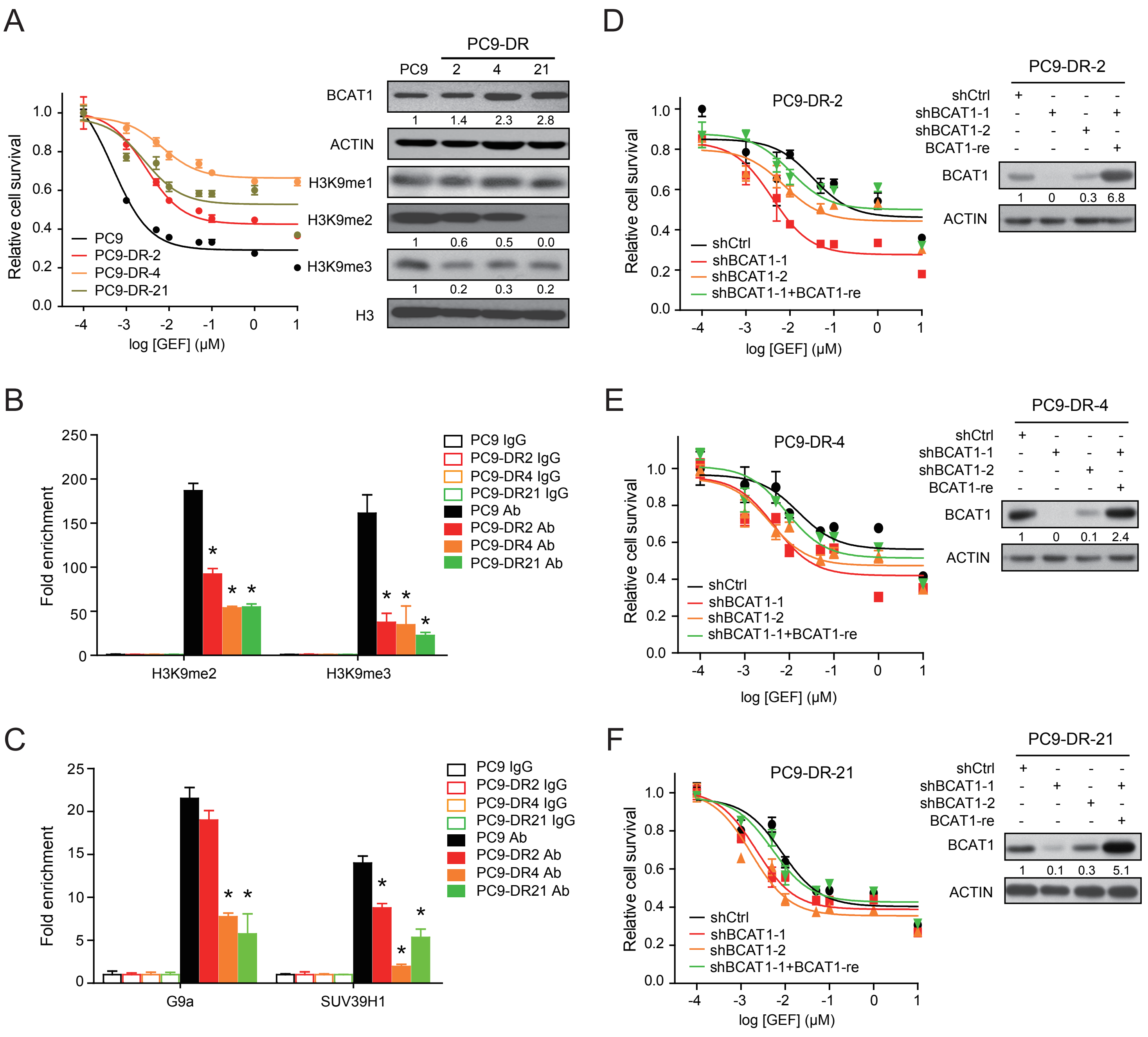
A
PC9-DR
PC9
2
4
21
BCAT1
1
1.4
2.3
2.8
ACTIN
H3K9me1
H3K9me2
1
0.6
0.5
0.0
H3K9me3
1
0.2
0.3
0.2
H3
Relative cell survival
PC9
PC9-DR-2
PC9-DR-4
PC9-DR-21
log [GEF] (μM)
B
Fold enrichment
H3K9me2
H3K9me3
PC9 IgG
PC9-DR2 IgG
PC9-DR4 IgG
PC9-DR21 IgG
PC9 Ab
PC9-DR2 Ab
PC9-DR4 Ab
PC9-DR21 Ab
C
Fold enrichment
G9a
SUV39H1
D
PC9-DR-2
shCtrl
shBCAT1-1
shBCAT1-2
shBCAT1-1+BCAT1-re
BCAT1-re
BCAT1
1
0
0.3
6.8
ACTIN
E
PC9-DR-4
1
0
0.1
2.4
F
PC9-DR-21
1
0.1
0.3
5.1

Figure S8

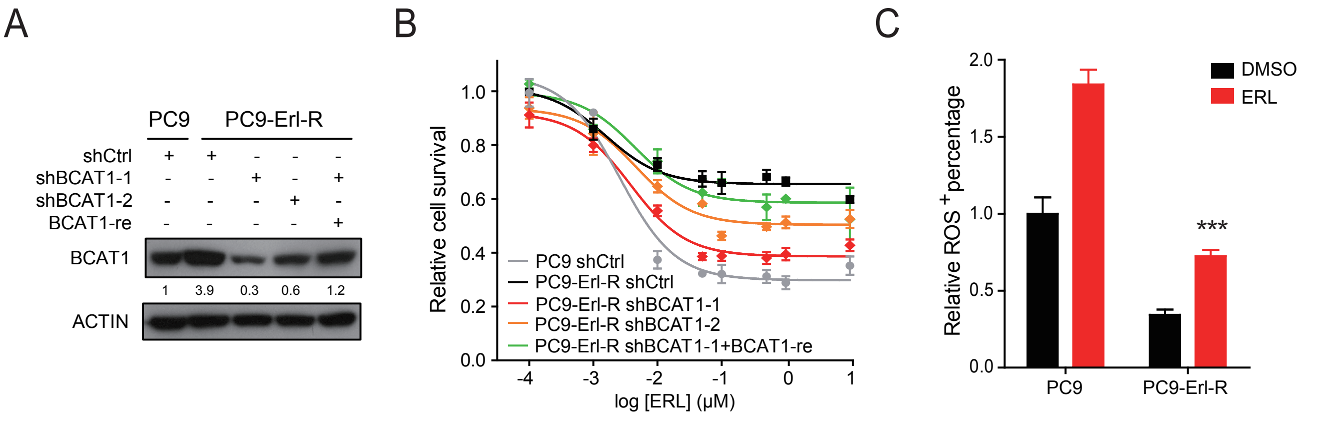
A
PC9
PC9-Erl-R
shCtrl
shBCAT1-1
shBCAT1-2
BCAT1-re
BCAT1
1
3.9
0.3
0.6
1.2
ACTIN
B
Relative cell survival
PC9 shCtrl
PC9-Erl-R shCtrl
PC9-Erl-R shBCAT1-1
PC9-Erl-R shBCAT1-2
PC9-Erl-R shBCAT1-1+BCAT1-re
log [ERL] (μM)
C
Relative ROS + percentage
DMSO
ERL
***
PC9
PC9-Erl-R

Figure S9

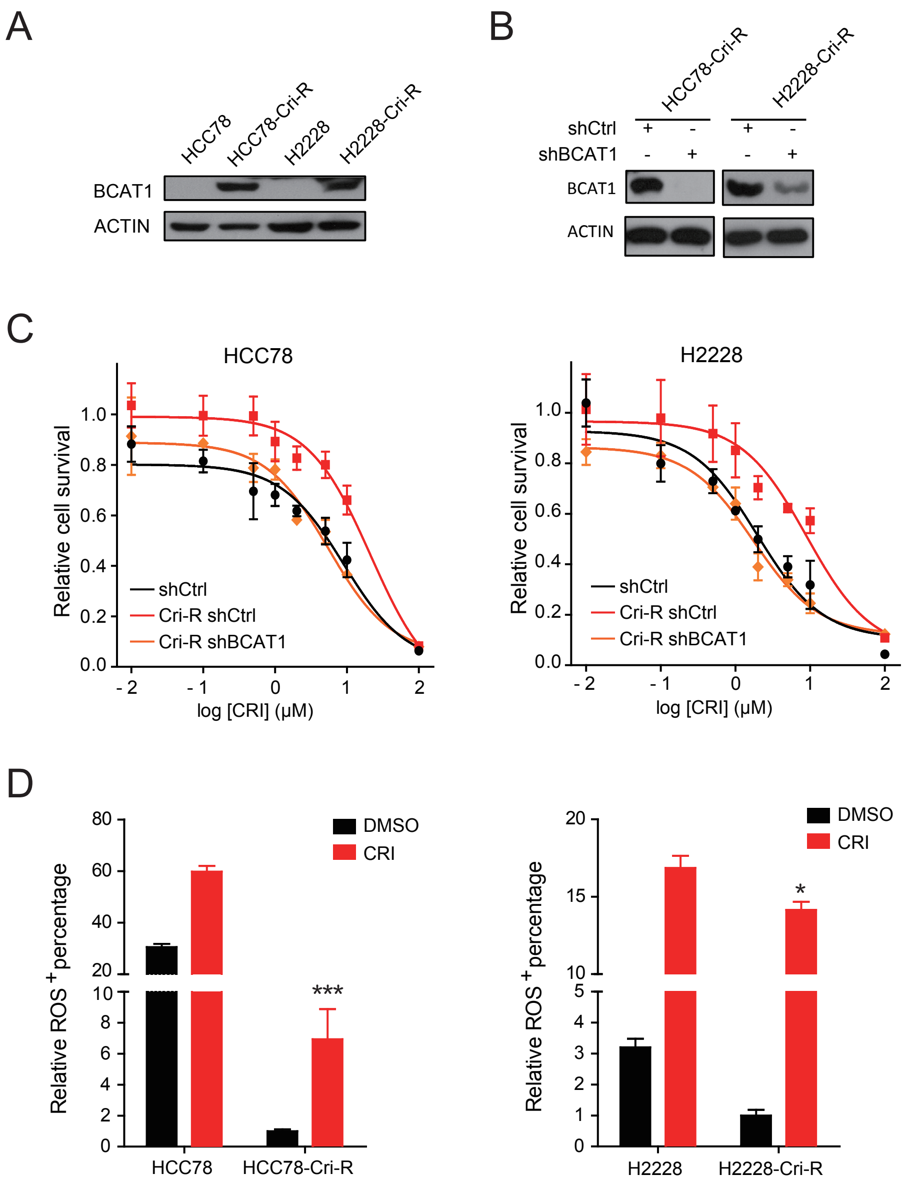
A
HCC78
HCC78-Cri-R
H2228
H2228-Cri-R
BCAT1
ACTIN
B
HCC78-Cri-R
H2228-Cri-R
shCtrl
shBCAT1
BCAT1
ACTIN
C
HCC78
H2228
Relative cell survival
shCtrl
Cri-R shCtrl
Cri-R shBCAT1
log [CRI] (μM)
D
Relative ROS + percentage
DMSO
CRI
***
*
HCC78
HCC78-Cri-R
H2228
H2228-Cri-R

Figure S10

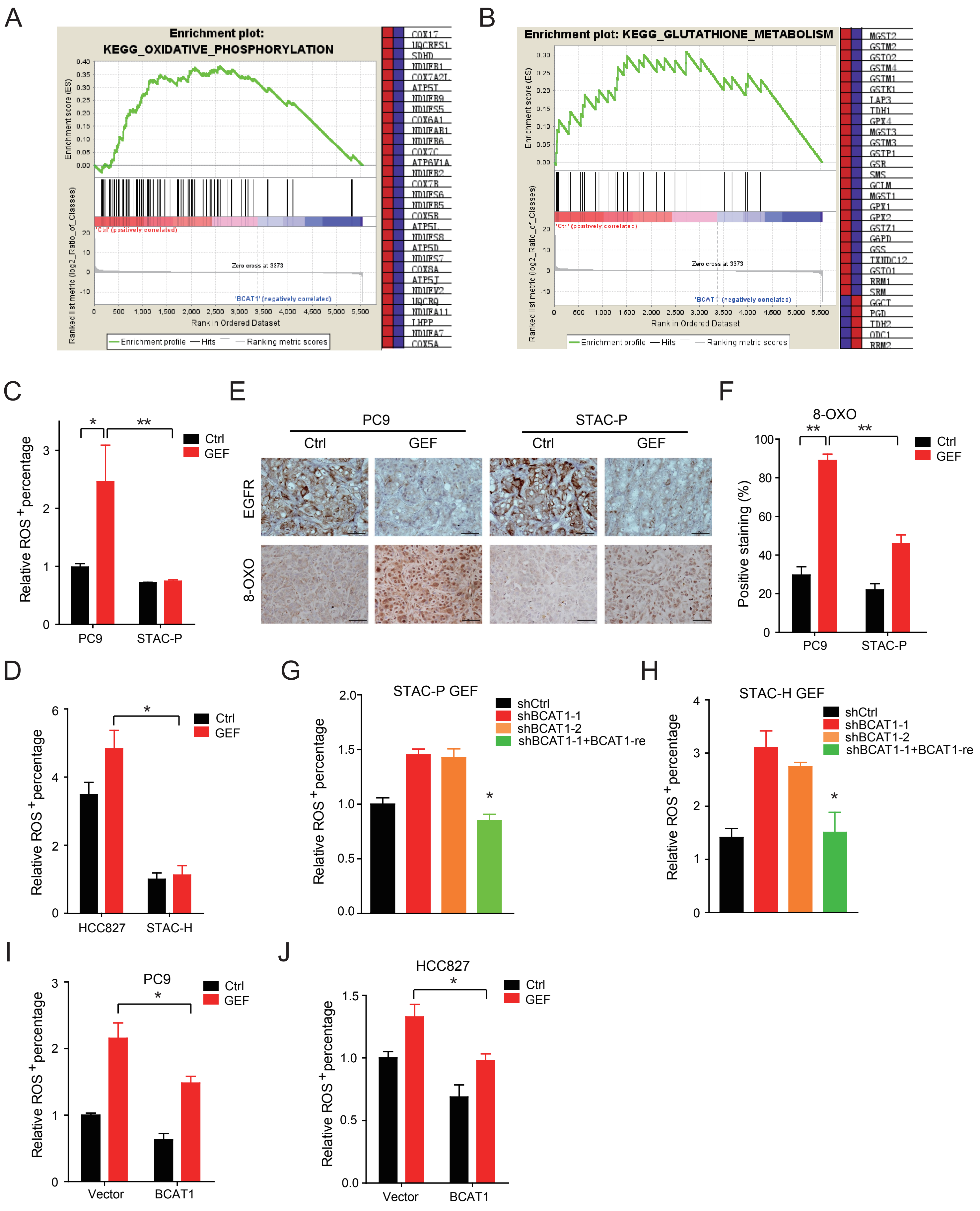
A
Enrichment plot: KEGG_OXIDATIVE_PHOSPHORYLATION
Enrichment score (ES)
Ranked list metric (log2_Ratio_of_Classes)
'Ctrl' (positively correlated)
Zero cross at 3373
'BCAT1' (negatively correlated)
Rank in Ordered Dataset
Enrichment profile
Hits
Ranking metric scores
B
Enrichment plot: KEGG_GLUTATHIONE_METABOLISM
C
Relative ROS+ percentage
Ctrl
GEF
PC9
STAC-P
E
PC9
STAC-P
Ctrl
GEF
EGFR
8-OXO
F
8-OXO
Positive staining (%)
D
HCC827
STAC-H
G
STAC-P GEF
shCtrl
shBCAT1-1
shBCAT1-2
shBCAT1-1+BCAT1-re
H
STAC-H GEF
I
PC9
Vector
BCAT1
J
HCC827

Figure S11

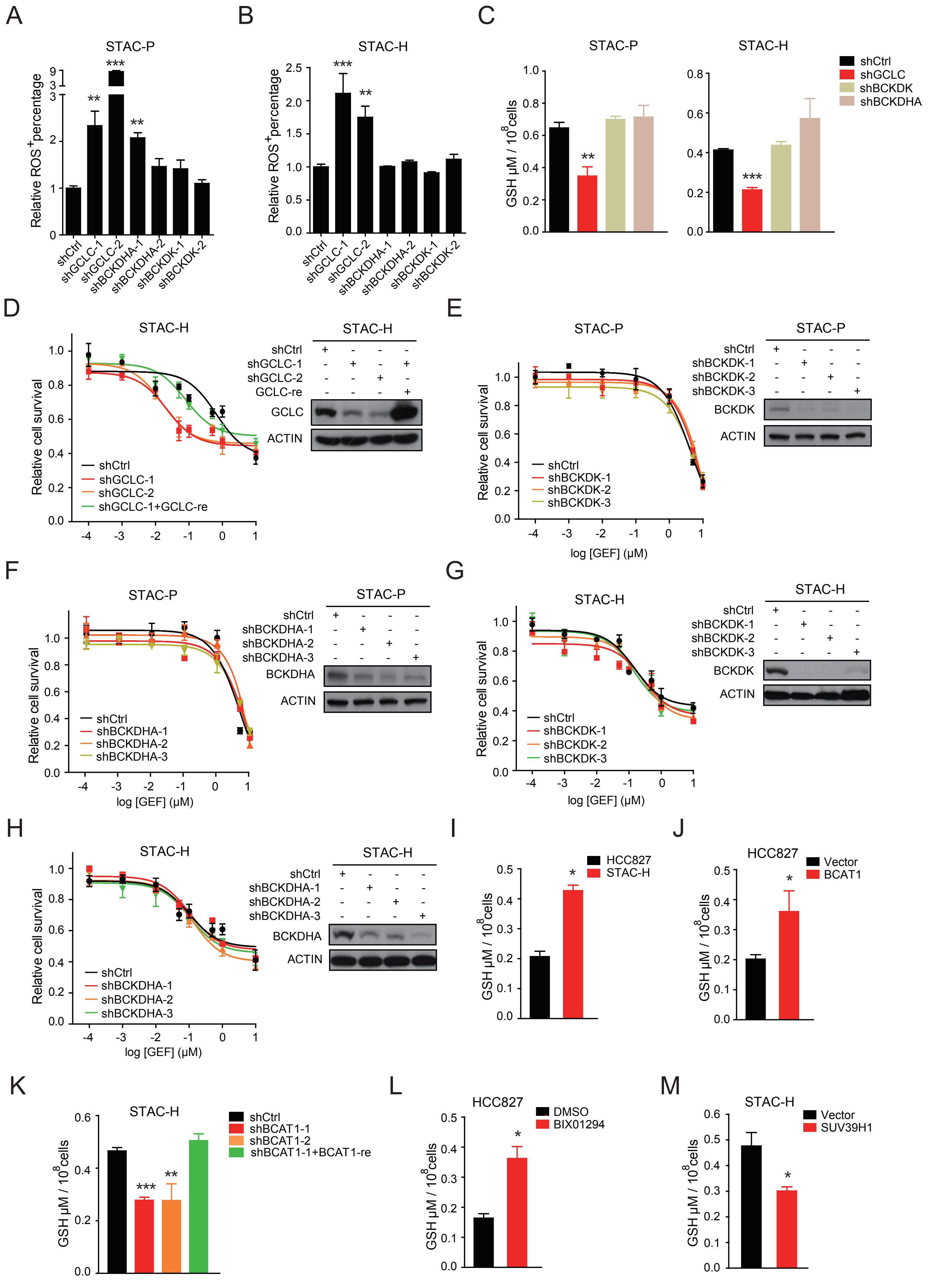
A
STAC-P
Relative ROS+percentage
shCtrl
shGCLC-1
shGCLC-2
shBCKDHA-1
shBCKDHA-2
shBCKDK-1
shBCKDK-2
B
STAC-H
C
STAC-P
STAC-H
GSH μM / 10^8cells
shCtrl
shGCLC
shBCKDK
shBCKDHA
D
STAC-H
Relative cell survival
shGCLC-1+GCLC-re
GCLC-re
GCLC
ACTIN
E
STAC-P
shBCKDK-3
BCKDK
log [GEF] (μM)
F
shBCKDHA-3
BCKDHA
G
H
I
HCC827
STAC-H
J
HCC827
Vector
BCAT1
K
STAC-H
shBCAT1-1
shBCAT1-2
shBCAT1-1+BCAT1-re
L
HCC827
DMSO
BIX01294
M
STAC-H
Vector
SUV39H1

Figure S12

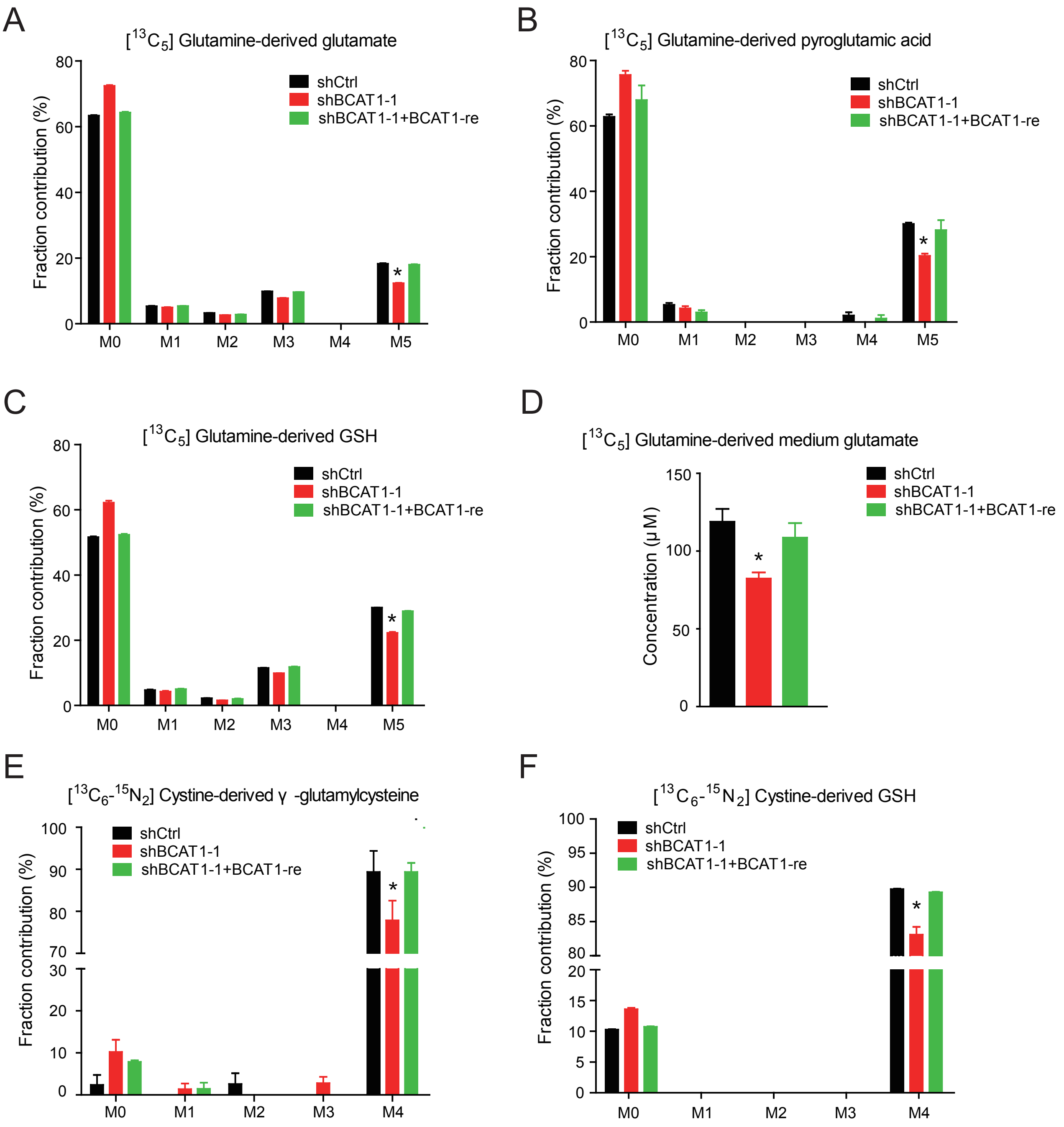
A
[13C5] Glutamine-derived glutamate
Fraction contribution (%)
shCtrl
shBCAT1-1
shBCAT1-1+BCAT1-re
M0
M1
M2
M3
M4
M5
B
[13C5] Glutamine-derived pyroglutamic acid
C
[13C5] Glutamine-derived GSH
D
[13C5] Glutamine-derived medium glutamate
Concentration (μM)
E
[13C6-15N2] Cystine-derived γ -glutamylcysteine
F
[13C6-15N2] Cystine-derived GSH

Figure S13

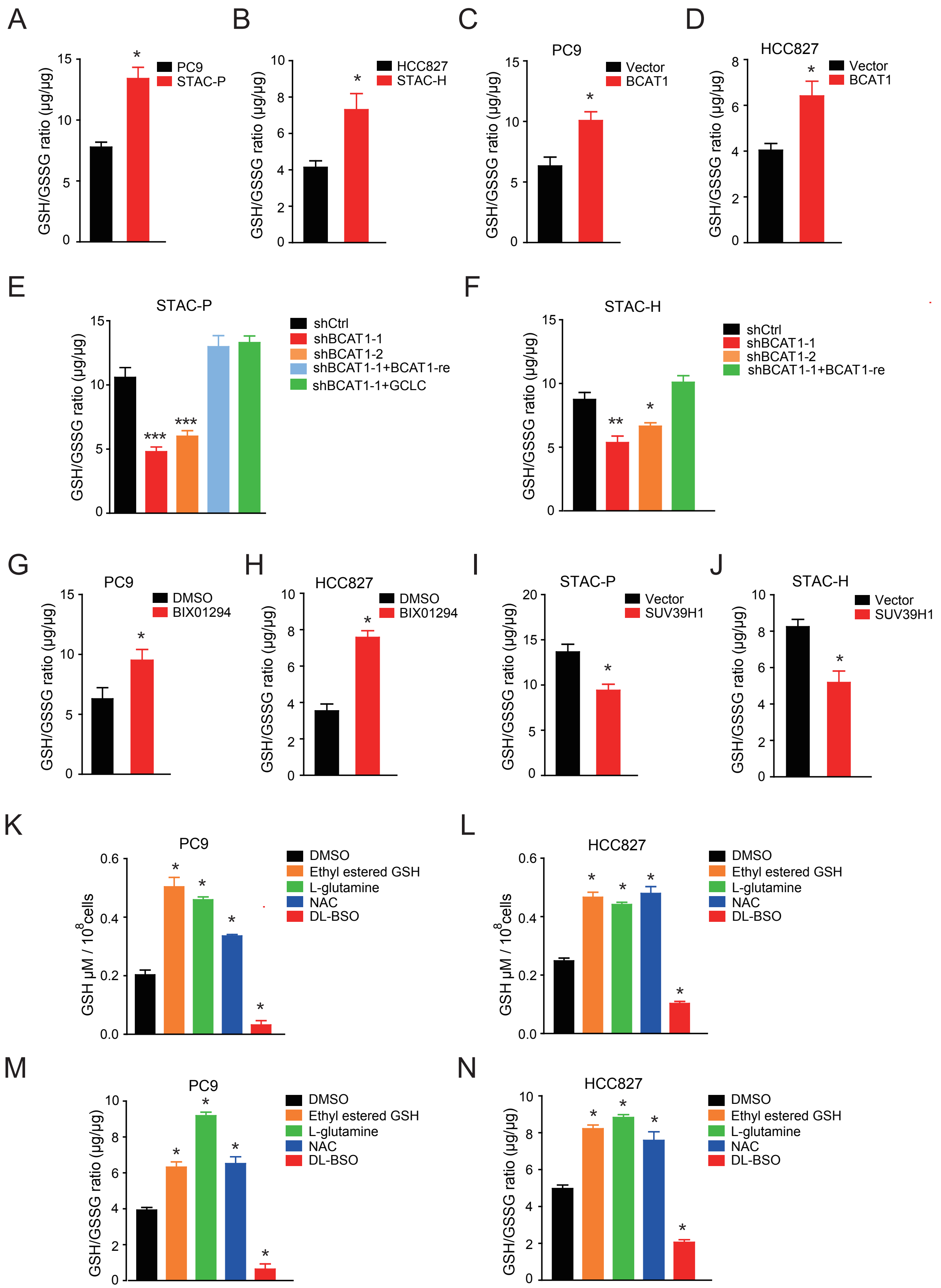

A
GSH/GSSG ratio (μg/μg)
PC9
STAC-P
B
HCC827
STAC-H
C
PC9
Vector
BCAT1
D
HCC827
Vector
BCAT1
E
STAC-P
shCtrl
shBCAT1-1
shBCAT1-2
shBCAT1-1+BCAT1-re
shBCAT1-1+GCLC
F
STAC-H
shCtrl
shBCAT1-1
shBCAT1-2
shBCAT1-1+BCAT1-re
G
PC9
DMSO
BIX01294
H
HCC827
DMSO
BIX01294
I
STAC-P
Vector
SUV39H1
J
STAC-H
Vector
SUV39H1
K
PC9
GSH μM / 10^8 cells
DMSO
Ethyl estered GSH
L-glutamine
NAC
DL-BSO
L
HCC827
M
PC9
N
HCC827


Figure S14

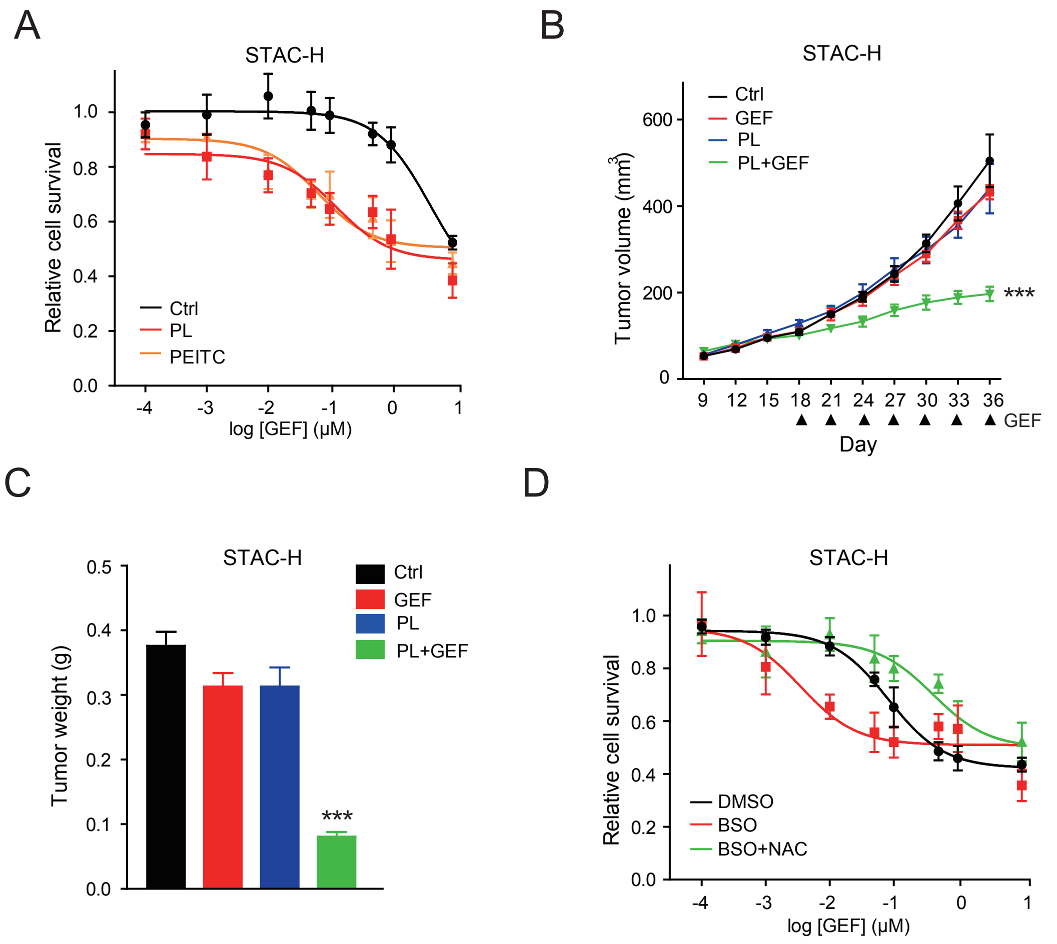
A
STAC-H
Relative cell survival
1.0
0.8
0.6
0.4
0.2
0.0
Ctrl
PL
PEITC
-4
-3
-2
-1
0
1
log [GEF] (μM)
B
STAC-H
Tumor volume (mm3)
600
400
200
0
Ctrl
GEF
PL
PL+GEF
***
9
12
15
18
21
24
27
30
33
36
GEF
Day
C
STAC-H
Tumor weight (g)
0.5
0.4
0.3
0.2
0.1
0.0
Ctrl
GEF
PL
PL+GEF
***
D
STAC-H
Relative cell survival
1.0
0.8
0.6
0.4
0.2
0.0
DMSO
BSO
BSO+NAC
-4
-3
-2
-1
0
1
log [GEF] (μM)

Figure S15

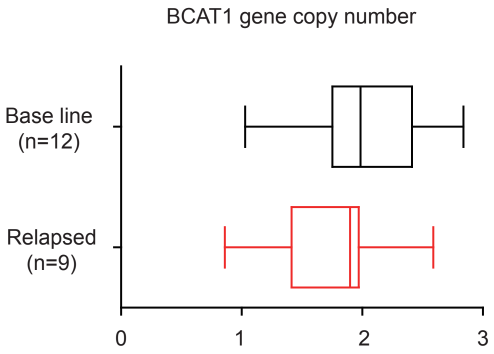


Figure S16

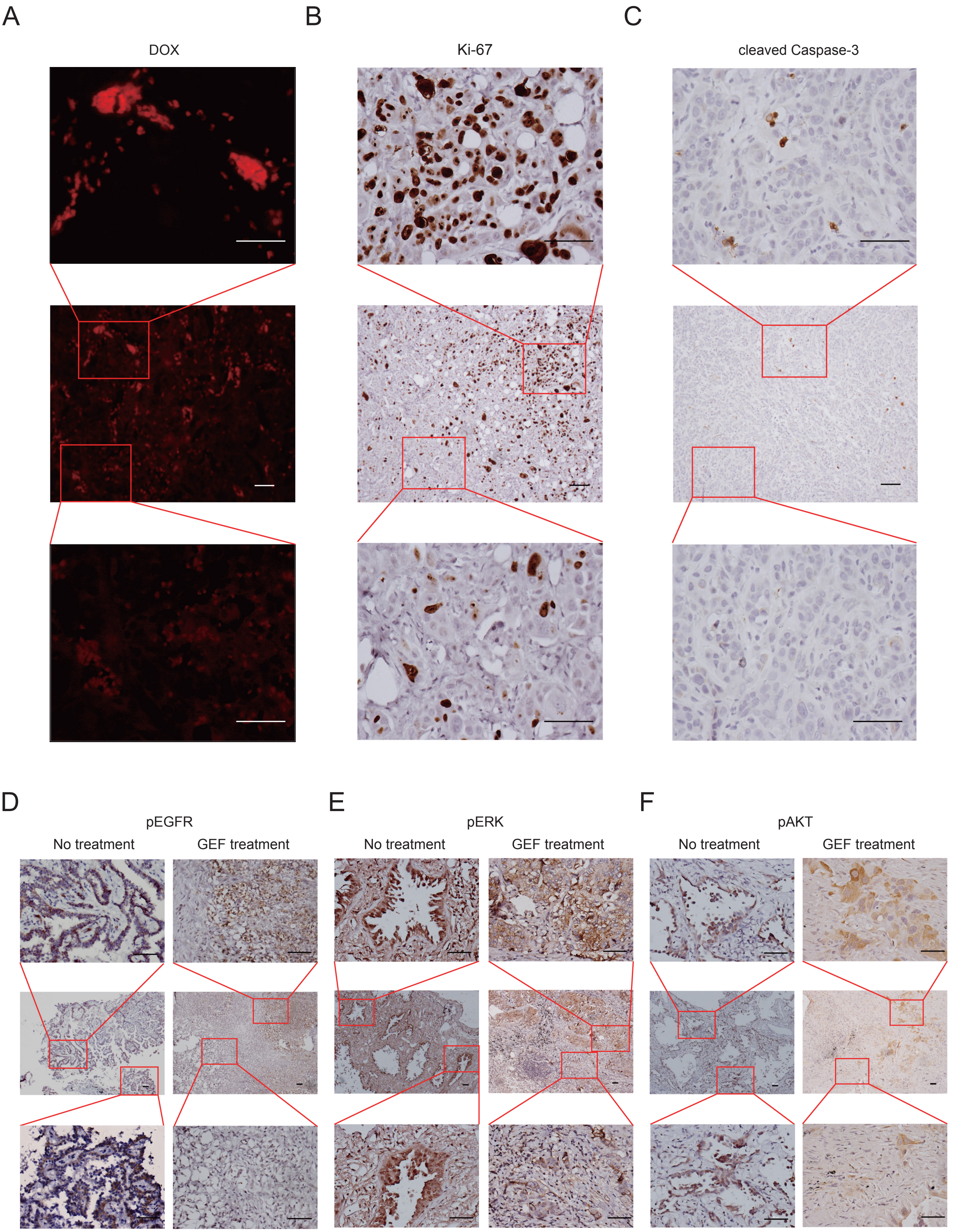
A
B
C
DOX
Ki-67
cleaved Caspase-3
D
E
F
pEGFR
pERK
pAKT
No treatment
GEF treatment
No treatment
GEF treatment
No treatment
GEF treatment

Figure S17

**STAR★METHODS**

Detailed methods are provided include the following:

**KEY RESOURCES TABLE**

| REAGENT | SOURCE | IDENTIFIER |
|---|---|---|
| **Antibodies** | | |
| pEGFR (Y1068) | CST | 2234 |
| pAKT (S473) | CST | 4060 |
| AKT | CST | 9272 |
| pERK1/2 (T202/Y204) | CST | 9106 |
| ERK | CST | 9102 |
| BCAT1 | CST | 12822 |
| H3K9me1 | ABclonal | A2358 |
| H3K9me2 | ABclonal | A2359 |
| H3K9me3 | ABclonal | A2360 |
| H3K9me2 | Abcam | AB1220 |
| H3K9me3 | Abcam | AB8898 |
| H3 | Proteintech | 17168-1-AP |
| ACTIN | Abcam | AB136452 |
| EGFR | ABclonal | A2909 |
| G9a | Proteintech | 11595-1-AP |
| G9a | Millipore | 07-551 |
| SUV39H1 | Proteintech | 10574-1-AP |
| SUV39H1 | Abcam | AB12405 |
| BCKDK | Sangon | D151861 |
| BCKDHA | Bioworld | BS711326 |
| GCLC | Proteintech | 12601-1-AP |
| 8-Oxoguanine | Abcam | AB48508 |
| Ki-67 | Leica Biosystems | NCL-Ki67p |
| IgG (mouse) | CST | 3900 |
| IgG (rabbit) | CST | 5415 |

| | | |
|---|---|---|
| H3K4me1 | A gift from Prof. Degui Chen (SIBCB) | N/A |
| H3K4me2 | A gift from Prof. Degui Chen (SIBCB) | N/A |
| H3K20me1 | A gift from Prof. Degui Chen (SIBCB) | N/A |
| H3K20me2 | A gift from Prof. Degui Chen (SIBCB) | N/A |
| H3K27me1 | A gift from Prof. Degui Chen (SIBCB) | N/A |
| H3K27me2 | A gift from Prof. Degui Chen (SIBCB) | N/A |
| H3K36me2 | A gift from Prof. Degui Chen (SIBCB) | N/A |
| H3K79me1 | A gift from Prof. Degui Chen (SIBCB) | N/A |
| **Recombinant DNA** | | |
| PCDH-CMV | This paper | N/A |
| PCDH-CMV-BCAT1 | This paper | N/A |
| PCDH-CMV-GCLC | This paper | N/A |
| PCDH-CMV-BCAT1-MUT | This paper | N/A |
| PCDH-CMV-GCLC-MUT | This paper | N/A |
| PCDH-CMV-G9a | This paper | N/A |
| PCDH-CMV-SUV39H1 | This paper | N/A |
| PLKO.1-U6-shluciferace | This paper | N/A |
| PLKO.1-U6-shBCAT1-1~3 | This paper | N/A |
| PLKO.1-U6-shGa9-1~2 | This paper | N/A |
| PLKO.1-U6-shSUV39H1-1~2 | This paper | N/A |
| PLKO.1-U6-shGCLC-1~2 | This paper | N/A |
| PLKO.1-U6-shBCKDK-1~2 | This paper | N/A |
| PLKO.1-U6-shBCKDHA-1~2 | This paper | N/A |
| psPAX2 | Addgene | 12260 |
| pMD2.G | Addgene | 12259 |
| **Chemicals** | | |
| Gefitinib | MCE | HY50895 |
| Erlortinib | SELLECK | S7786 |

| | | |
|---|---|---|
| BIX01294 | SELLECK | S8006 |
| Cisplatin | SELLECK | S1166 |
| N-Acetyl-L-cystine (NAC) | MCE | HY-B0215 |
| L-Glutamine | MCE | HY-N0390 |
| Piperlongumine (PL) | SELLECK | S7551 |
| Phenethyl isothiocyanate (PEITC) | Sigma | 253731 |
| DL-Buthionine-sulfoximine (BSO) | Sigma | 19176 |
| GSH-ethyl ester | Cayman | 14953 |
| **Cell lines** | | |
| PC9 | ATCC | N/A |
| HCC827 | ATCC | N/A |
| SH450 | (Zheng et al., 2011) | N/A |
| HEK293T | ATCC | N/A |
| **Mice** | | |
| Nude mice | SLAC | N/A |
| NOD-SCID mice | SLAC | N/A |
| **Critical Commercial Assays** | | |
| Reactive Oxygen Species Assay Kit | Beyotime | S0033 |
| Glutathione Fluorometric Assay Kit | BioVision | K264-100 |
| **Software and Algorithms** | | |
| GraphPad prism | GraphPad Software | https://www.graphpad.com/ |
| ImageJ software | Open source | https://imagej.nih.gov/ij/ |

### CONTACT FOR REAGENT AND RESOURCE SHARING

### EXPERIMENTAL MODEL AND SUBJECT DETAILS

#### Cell Culture Studies

PC9, HCC827, HCC78 and H2228 cells were purchased from ATCC and were free of mycoplasma contamination. Cells were grown in DMEM (Hyclone) with 8% FBS (Gibco). SH450 cells were established as described previously (Zheng et al., 2011) and were grown in DMEM (Hyclone) with 8% FBS (Gibco).

**Animal Studies**

Mice were housed in a specific pathogen-free environment at the Shanghai Institute of Biochemistry and Cell Biology and treated in strict accordance with protocols approved by the Institutional Animal Care and Use Committee of the Shanghai Institutes for Biological Sciences, Chinese Academy of Sciences. Nude mice at 6 weeks old were subcutaneously transplanted with $5x10^6$ cells until palpable tumors formed. The mice were then randomly divided into different groups (in each group n=6) and treated with 50 mg/kg GEF or 3.125 mg/kg GEF alone or in combination with 3 mg/kg PL or 450 mg/kg BSO with indicated administrated frequency via intraperitoneal injection. NAC was prepared in the drinking water with the final 40mM concentration for the treatment group during the experiment (Anastasiou et al., 2011). Tumors were monitored every other day and tumor volume was calculated as follows: tumor volume ($mm^3$) = (length x $width^2$) / 2. Relative tumor growth was calculated by normalizing the tumor volumes from different days to the starting volume of individual tumor.

**Human Samples**

This study is approved by the institutional review committees of Shanghai pulmonary hospital, Tongji University. Patients gave written informed consents. A total of 119 biopsy specimens including 80 samples before EGFR TKI treatment as the baseline and 39 samples with acquired EGFR TKI resistance were used for clinical relevance analyses, among which 4 were matched samples from single patients. The 39 specimens with acquired resistance to EGFR TKIs were also analyzed for T790M mutation, MET amplification, HER2 amplification as well as SCLC transformation. Immunostaining of H3K9me2, 8-OXO, BCAT1 were blindly scored. As for the correlation analyses, the specimens were classified into three groups based on the IHC score of H3K9me2, 8-OXO, BCAT1 from low (IHC score: 0-100), medium (IHC score: 101-200) to high (IHC score: 201-300) and the sample numbers of each group were then calculated.

### METHOD DETAILS

#### Cell Proliferation Assay

Cells were seeded in sextuplicate in 96-well plates and stained with 3-(4,5-dimethyl-2-thiazolyl)-2,5-diphenyl-2-H-tetrazolium bromide (MTT) and assessed with Epoch multi-volume spectrophotometer system (570nm/630nm) at indicated time points. The survival ratio of drug-treated cells was calculated by dividing the fluorescence obtained from the drug-treated cells by the fluorescence obtained from the control (no drug)-treated cells. Experiments were done in quadruplicate. Data were shown as mean ± SEM.

#### Cell Survival Assay

Cells were seeded in triplicate in 96-well plate overnight and given indicated doses of various compounds 24 hrs later and continuously cultured for another 72 hrs. Cells were then stained with MTT and assessed with Epoch multi-volume spectrophotometer system (570nm/630nm). The growth ratio was calculated by normalizing the fluorescence obtained to the fluorescence of the first day.

#### TKI Pre-treatment Experiments

PC9, HCC827 or SH450 cells were pretreated with GEF or ERL (10nM) for 2 hrs and then replaced with complete medium without TKI for 0, 2, 4, 6 hrs. Cells were then treated with serial doses of GEF or ERL (50, 100, 500, 1000nM) for 0.5 hr and then cultured in complete medium without TKI for 72 hrs before subjected to MTT assay. For three cycles of pre-treatment experiments, cancer cells were pre-treated with 10nM TKI for three cycles (each cycle containing 2 hrs treatment followed by 2 hrs recovery). Experiments were done in quadruplicate. Data were shown as mean ± SEM.

#### Generation of Sublethal TKI Adapted Cells (STAC)

PC9 or HCC827 or SH450 cells were continuously treated with GEF or ERL (10nM) for over three months. Cells were passaged every three days and the half-maximal dosage effect (IC50) values were measured every four passages until the drug resistance was acquired. Cells at different passages were also collected for protein and RNA extractions and subjected to biochemical analyses. After the drug resistance development, the Sublethal TKI Adapted Cells (STACs) were cultured in complete medium without TKI for over 70 passages and the IC50 were measured intermittently.

**Plasmid Construction and Virus Infection**

Human G9a, SUV39H1 and BCAT1 cDNAs were amplified through PCR using the cDNA from normal human cell line 293T cells as the template. The primers used for these PCR were described in Supplementary Materials and Methods. All cDNAs were cloned into the lentiviral vector PCDH-CMV with a puromycin-resistance gene. For knockdown experiments, shRNA constructs from Generay Company were used and the sequences were described in Supplementary Materials and Methods. All shRNAs were cloned into the pLKO.1 lentiviral vector with a puromycin-selection marker as described previously (Li et al., 2015). Empty pCDH and/or pLKO.1 vector were used as controls. Lentiviral delivery of ectopic expression plasmid and shRNAs directed against genes were performed as described previously (Gao et al., 2014). Plasmids were packaged into lentiviral particles by co-transfection with packaging plasmids into HEK-293T cells and the filtered cell culture supernatant was then used to infect cells.

**Gene Amplification Analyses**

Total cell genomic DNAs were prepared as described previously (Gao et al., 2010) and were further used for amplification detection. Real-time PCR was performed using ABI7500 sequence detection system (Perkin Elmer Life Sciences, Shelton, CT). The list of primers was showed in the supplemental materials.

**Real-time PCR Analyses**

Total RNAs were prepared as described previously (Gao et al., 2010) and retro-transcribed into first-strand cDNA using the first-strand synthesis system following the manufacturer’s protocol (Invitrogen, Carlsbad, CA). Real-time PCR was performed using ABI7500 sequence detection system (Perkin Elmer Life Sciences, Shelton, CT). The sequences of primers were listed in the supplemental materials.

Ectopic expression and knockdown efficiency were also analyzed by real-time PCR through normalization to the control group. Experiments were done in quadruplicate. Data are presented as mean ± SEM.

#### Western Blot Assay

Cells lysates were prepared and subjected to western blot analysis as described previously (Gao et al., 2010) with following primary antibodies: pEGFR (CST-2234, Cell Signaling Technologies), EGFR (A2909, ABclonal), pAKT (CST-4060, Cell Signaling Technologies), AKT (CST-9272, Cell Signaling Technologies), pERK (CST-9106, Cell Signaling Technologies), ERK (CST-9102, Cell Signaling Technologies), BCAT1 (CST-12822, Cell Signaling Technologies), ACTIN (AB136452, Abcam), H3K9me1 (A2358, ABclonal), H3K9me2 (A2359, ABclonal), H3K9me3 (A2360, ABclonal), G9a (11595-1-AP, Proteintech), SUV39H1 (10574-1-AP, Proteintech), BCKDK (D151861, Sangon), BCKDHA (BS711326, Bioworld), GCLC (12601-1-AP, Proteintech) and H3 (17168-1-AP, Proteintech). Antibodies of H3K4Me1, H3K4Me2, H3K20Me1, H3K20Me2, H3K27Me1, H3K27Me2, H3K27Me3, H3K36Me2 and H3K79Me1 were gifts from Prof. Degui Chen. Image J was used to calculate the strength of bands.

#### Immunohistochemical Staining

Immunohistochemical (IHC) staining was performed as described previously (Li et al., 2015). Antibodies against the 8-Oxoguanine (AB48508, Abcam), Ki-67 (NCL-Ki67p, Leica Biosystems), H3K9me2 (AB1220, Abcam) and BCAT1 (CST-12822, Cell Signaling Technologies) were used. The IHC staining was blindly scored and the IHC score was calculated as previously described (Pirker et al., 2012). Briefly, for sections with analyzable tumor cells, staining intensity was scored in four categories: no staining (0), weak positive staining (1), intermediate positive staining (2) and strong positive staining (3). We used the prospectively collected IHC data to generate IHC scores on a scale of 0–300. By integration of the data relating to the intensity and frequency of staining, the IHC score was calculated with the formula: percentage of cells staining x positive staining strength.

**H3K9 Methyltransferase Enzyme Activity Test**

Cells were seeded for 24 hrs before collected. The protein of nuclear was extracted by using Nuclear and Cytoplasmic Protein Extraction Kit (P0027, Beyotime) according to the manufacturer's instructions. And the protein was stored at -20°C. The enzyme activity of HMT for H3K9 was detected by using EpiQuik Histone Methyltransferase Activity Assay Kit (P-3003, EpiQuik) according to the manufacturer's instructions.

**Chromatin Immunoprecipitation Assay**

Cells were cross-linked with 1% formaldehyde for 5 min at room temperature, lysed by SDS lysis buffer and sonicated to generate DNA fragments with an average size of 500bp. After pre-clearing with Protein A/G beads, antibody against H3K9me2 (AB1220, Abcam), H3K9me3 (AB8898, Abcam), G9a (07-551, Millipore), SUV39H1 (AB12405, Abcam), rabbit IgG (CST-3900, Cell Signaling Technologies) or mouse IgG (CST-5415, Cell Signaling Technologies) was added to cell lysate and incubated at 4$^{o}$C overnight. DNA cross-linked with antibodies was then pulled down with Protein A/G beads, washed, and purified with MinElute PCR purification kit (NO.28004, QIANGEN). Aliquots of ChIP-enriched DNA and whole-cell lysate DNA were subjected to real-time PCR analyses.

**GSH Density Measurement**

GSH concentration was measured using the Glutathione Fluorometric Assay Kit (K264-100, Biovision). Cells are seeded and treated with various compounds for 24 hr. $4 \times 10^6$ cells were then homogenized with ice-cold Glutathione Assay Buffer. The homogenate was mixed with PCA evenly and spun down for supernatant collection. Ice-cold KOH was then added to PCA-preserved samples to precipitate PCA. After spinning, neutralized samples and standard samples were transferred to a 96-well plate, added with OPA Probe into each well, and incubated at room temperature for 40 min. For GSSG, GSH quencher was added to an equal volume of neutralized sample for GSH mensuration, after RT incubation, add Reducing Agent Mix to destroy the excess GSH Quencher and convert GSSG to GSH and then the mixed buffer was subjected to the next step. Samples and standards were measured using a fluorescence plate reader (Ex/Em = 340/420 nm).

**Intracellular Reactive Oxygen Species (ROS) Detection**

Intracellular oxidative stress was assayed by measuring intracellular oxidation of 2, 7 -dichlorofluorescin (DCFH) (S0033, Beyotime). Cells were incubated with DCFH Probes for 30 min. And fluorescence of the oxidized form of DCFH (DCF) was measured using a flow cytometer LSRⅡ (Becton Dickenson) as previously reported (Li et al., 2015).

**UHPLC-qTOF-MS Analysis**

For metabolite extraction, the cell samples were corrected to ~5×$10^6$ cells per sample in 2-mL Eppendorf tubes with 1.6 mL of 80% (v/v) methanol solution for each sample. Each sample was centrifuged at 14,000 g at 4 ºC and the supernatant was evaporated to dryness at 20 ºC. The residue was reconstituted by adding 300 μL of 75% (v/v) acetonitrile solution containing 5 mM ammonium formate, and then vortexed for 30 s before centrifugation at 14,000 g for 10 min at 4°C. For each sample, 1μL of the supernatant was injected into the LC/MS for measurement. The quality control (QC) sample was a mixture prepared from an equal amount in each sample supernatant and analyzed with the same procedure as that for the experimental samples. The liquid chromatography with Agilent uHPLC (Binary Pump, Multisampler and Column Comp) was prepared and all chromatographic separations were performed with a Waters BEH Amide column (100mm × 2.1mm, 1.7μm). The mobile phase consisted of ammonium hydroxide and 10 mM $NH_4Ac$ in LC-MS grade water (mobile phase A) and LCMS grade acetonitrile (mobile phase B) run at a flow rate of 0.3 mL/min. The gradient program was initiated at 90% B held for 1 min, decreased to 60% B in 9 min and held for 5 min, and the post time was set 7 min. Positive ion mode was utilized in the mass spectrometer with a 4.0kV capillary voltage. Nozzle voltage 750v and scan range from 50-1100 Da were applied. The source gas and sheath gas temperature was set 325°C and 325°C, respectively.

**Quantification of [$^{13}C_5$]-Glutamate in Condition Medium**

A 20μL thawed medium, 80 μL methanol was added. After votex and centrifugation at 4 °C and 14000 g for 15 min, 50 μL supernatant was took and mixed with 50 μL internal standard (100ng/mL [$^{13}C_6$,$^{15}N$]-leucine) and 100 μL 50% acetonitrile prior to UPLCMS/MS analysis.The UPLC-MS/MS analysis was performed on a Waters Acquity UPLC system (Waters, Milford, MA) coupled to a Triple Quad™ 6500plus tandem mass spectrometer (AB Sciex, Framingham, MA), and 2μL sample was injected onto a Waters BEH Amide column (100mm × 2.1mm, 1.7μm) at a flow rate of 0.3 mL/min. The mobile phase consisted of (A) water and (B) 90% acetonitrile, both with 10 mM $NH_4Ac$ and 0.2% formic acid. The chromatographic separation was conducted by a gradient elution program as follows: 0 min, 90% B and held to 0.5 min; 3.5 min, 70%B; 6 min, 40% B; 6.5 min, 40% B; 6.6 min, 90% B and held to 8.5 min. The column temperature was 40 ºC. The analytes eluted from column were ionized in an electro spray ionization source in positive mode (ESI+). Source temperature: 500 °C, curtain gas (CUR): 35 psi, ion source gas 1 (GS1): 50 psi, ion source gas 2 (GS2): 60 psi, collision gas (CAD): 8 psi, ion spray voltage (IS): 5500 V, entrance potential (EP): 10 eV, collision cell exit potential (CXP1): 9 eV. The multiple reaction monitoring (MRM) was used to acquire data in optimized MRM transition (precursor >product), declustering potential (DP), and collision energy (CE). The MRM transition, DP, and CE were set as 153>88, 40V, and 38 V for [$^{13}C_5$]-glutamate, while 139>92, 50V, and 27V for [$^{13}C_6$,$^{15}N$]-leucine. AB Sciex Analyst software (version 1.5.2) was used to control instruments, acquire and analyze data.

**Bioinformatics Analysis**

Total RNAs from PC9 and STAC-P cells were used for microarray and RNA-seq experiments. For microarray data analyses, we normalized the expression with custom CDF of Refseq annotation and considered a gene with over 2-fold change as differentially expressed. For RNA-seq data, we aligned the reads to hg19 using TopHat2 and gene counts were calculated via HTSeq-count tools. We performed differentially expressed genes analyses using DESEQ in R. We also compared the different gene sets correlation between arrays and sequencing data and found that these two datasets had Pearson correlation of 0.428 with significant p-value. We then picked up the upregulated genes in both arrays and sequencing data and found 22 upregulated candidates in total.

Total RNAs extracted from STAC-P with or without BCAT1 knockdown were also used for RNA-seq analysis. For this, we calculated the fold change of each probe (a gene with over 2-fold change was considered as differentially expressed) and then used GSEA to rank the probes and analyzed the enrichment based on Hypergeometric tests as described previously (Subramanian et al., 2005). The heat map of the signature genes was generated in R by the function heatmap.2 based on their expression levels. The permutation type was gene set. Classic enrichment statistic was used to find the enriched genes. Log2_Ratio_of_Classes was used as metric for ranking genes. All the array and sequencing data have been deposited on the GEO database with the accession codes GSE79433, GSE79763 and GSE79764.

**PDX Model Establishment**

The tumor collection and transplantation were approved by IRB and patients gave written informed consents. Human lung tumors were freshly dissected and subcutaneously transplanted into NOD-SCID mice from SLAC to establish PDX model. The PDX model at passage 3 was used for the treatment experiment. Nude mice at 6 weeks old were subcutaneously transplanted with PDX tumors (~50mm$^3$) until palpable tumors formed. The mice were then randomly divided into different groups (in each group n=6) and treated with 50mg/kg GEF alone or in combination with 3 mg/kg PL or 450 mg/kg BSO with indicated administrated frequency via intraperitoneal injection. NAC was prepared in the drinking water with the final 40mM concentration for the treatment group during the experiment (Anastasiou et al., 2011). Tumors were monitored every other day or every day and tumor volume was calculated as follows: tumor volume (mm$^3$) = (length x width$^2$) / 2. Relative tumor growth was calculated by normalizing the tumor volumes from different days to the starting volume of individual tumor.

**Clinical Character Definition**

Objective tumor response was determined using the response evaluation criteria in solid tumors (RECIST Version 1.1). The patients had computed tomography scan covering target lesions 4-6 weeks later after the initiating of EGFR TKI treatment and then every 8 weeks or when the symptom indicated. Brain or bone metastases were evaluated every 6 months by magnetic resonance imaging or bone scintigraphy. According to the Response Evaluation Croteria in Solid Tumors guideline, response to the therapy was categorized into four groups: complete response, partial response, stable disease and progression disease. Progression-free survival (PFS) was calculated from the date of the beginning of EGFR TKI treatment to the date of tumor progression or death. Overall survival (OS) was calculated as the time from the beginning of therapy to death or last follow-up. PD (progressed disease), SD (stabled disease) and PR (partially response) were used to classify EGFR TKI clinical response.

**Quantification and Statistical Analysis**

Data were presented as mean ± SEM unless specified. The statistical significance of differences was determined using the unpaired Student's t test, Mann-Whitney U test, or the One-way ANOVA. Kendall's tau correlation test was used to analyze gene expression correlation. Kaplan-Meier analysis with log-rank test was used to compare PFS between subgroups. All statistical analyses were carried out using SPSS 16.0 or GraphPad Prism 5 software (San Diego, CA), and $p$ value $< 0.05$ was considered to be statistically significant.

### Primer List

| | |
|---|---|
| H-shGCLC-F-1 | CCGG GTCATCAATGTACCAATATTT CTCGAG AAATATTGGTACATTGATGACTTTTTG |
| H-shGCLC-R-1 | AATTCAAAAA GTCATCAATGTACCAATATTT CTCGAG AAATATTGGTACATTGATGAC |
| H-shGCLC-F-2 | CCGG GCCATTGAAGAACAATAACTA CTCGAG TAGTTATTGTTCTTCAATGGCTTTTTG |
| H-shGCLC-R-2 | AATTCAAAAA GCCATTGAAGAACAATAACTA CTCGAGTAGTTATTGTTCTTCAATGGC |
| H-shBCKDHA-F-1 | CCGG GCTGAAGCTCTACAAGAGCAT CTCGAG ATGCTCTTGTAGAGCTTCAGCTTTTTG |
| H-shBCKDHA-R-1 | AATTCAAAAA GCTGAAGCTCTACAAGAGCAT CTCGAGATGCTCTTGTAGAGCTTCAGC |
| H-shBCKDHA-F-2 | CCGG CCTACTCTTCTCAGACGTGTA CTCGAG TACACGTCTGAGAAGAGTAGGTTTTTG |
| H-shBCKDHA-R-2 | AATTCAAAAA CCTACTCTTCTCAGACGTGTA CTCGAGTACACGTCTGAGAAGAGTAGG |
| H-shBCKDHA-F-3 | CCGGCCTACTCTTCTCAGACGTGTACTCGAGTACACGTCTGAGAAGAGTAGGTTTTTG |
| H-shBCKDHA-R-3 | AATTCAAAAACCTACTCTTCTCAGACGTGTACTCGAGTACACGTCTGAGAAGAGTAGG |
| H-shBCKDK-F-1 | CCGG GATCTGATCATCAGGATCTCA CTCGAGTGAGATCCTGATGATCAGATCTTTTTG |
| H-shBCKDK-R-1 | AATTCAAAAA GATCTGATCATCAGGATCTCA CTCGAGTGAGATCCTGATGATCAGATC |
| H-shBCKDK-F-2 | CCGG TCAGGACCCATGCACGGCTTT CTCGAG AAAGCCGTGCATGGGTCCTGATTTTTG |
| H-shBCKDK-R-2 | AATTCAAAAA TCAGGACCCATGCACGGCTTTCTCGAGAAAGCCGTGCATGGGTCCTGA |
| H-shBCKDK-F-3 | CCGGCCAGCACCAGTTCCGTCATTCCTCGAGGAATGACGGAACTGGTGCTGGTTTTTG |
| H-shBCKDK-R-3 | AATTCAAAAACCAGCACCAGTTCCGTCATTCCTCGAGGAATGACGGAACTGGTGCTGG |
| H-shBCAT1-1-F | CCGG CCCAATGTGAAGCAGTAGATA CTCGAGTATCTACTGCTTCACATTGGG TTTTTG |
| H-shBCAT1-1-R | AATTCAAAAA CCCAATGTGAAGCAGTAGATA CTCGAG TATCTACTGCTTCACATTGGG |
| H-shBCAT1-2-F | CCGG CCATTCTTCAAGACTTAGTTA CTCGAGTAACTAAGTCTTGAAGAATGG TTTTTG |
| H-shBCAT1-2-R | AATTCAAAAA CCATTCTTCAAGACTTAGTTACTCGAGTAACTAAGTCTTGAAGAATGG |
| H-shBMP5-1F | CCGGGCTGCCGGACAGATATATATTCTCGAGAATATATATCTGTCCGGCAGCTTTTTG |
| H-shBMP5-1R | AATTCAAAAAGCTGCCGGACAGATATATATTCTCGAGAATATATATCTGTCCGGCAGC |
| H-shBMP5-2F | CCGGCCACGAAAGACGGGAAATACACTCGAGTGTATTTCCCGTCTTTCGTGGTTTTTG |
| H-shBMP5-2R | AATTCAAAAACCACGAAAGACGGGAAATACACTCGAGTGTATTTCCCGTCTTTCGTGG |
| H-shBMP5-3F | CCGGGAGTCGGAGTACTCAGTAAGGCTCGAGCCTTACTGAGTACTCCGACTCTTTTTG |

| | |
|---|---|
| H-shBMP5-3R | AATTCAAAAAGAGTCGGAGTACTCAGTAAGGCTCGAGCCTTACTGAGTACTCCGACTC |
| H-shFAM171A1-F-1 | CCGGTATGAAGATGTCGTCCAAATACTCGAGTATTTGGACGACATCTTCATATTTTTG |
| H-shFAM171A1-R-1 | AATTCAAAAATATGAAGATGTCGTCCAAATACTCGAGTATTTGGACGACATCTTCATA |
| H-shFAM171A1-F-2 | CCGG TTGCGAGGATTAGACGGAAAT CTCGAGATTTCCGTCTAATCCTCGCAATTTTTG |
| H-shFAM171A1-R-2 | AATTCAAAAA TTGCGAGGATTAGACGGAAAT CTCGAGATTTCCGTCTAATCCTCGCAA |
| H-shCYB5R2-F-1 | CCGGCAGAGGCTTTGTGGACCTAATCTCGAGATTAGGTCCACAAAGCCTCTGTTTTTG |
| H-shCYB5R2-R-1 | AATTCAAAAACAGAGGCTTTGTGGACCTAATCTCGAGATTAGGTCCACAAAGCCTCTG |
| H-shHSD3B1-F-1 | CCGGGAAGGTTTCTGTCCTAATCATCTCGAGATGATTAGGACAGAAACCTTCTTTTTG |
| H-shHSD3B1-R-1 | AATTCAAAAAGAAGGTTTCTGTCCTAATCATCTCGAGATGATTAGGACAGAAACCTTC |
| H-sh4-Mar-F-1 | CCGGCCTCCTCAGATGACTTCTGTACTCGAGTACAGAAGTCATCTGAGGAGGTTTTTG |
| H-sh4-Mar-R-1 | AATTCAAAAACCTCCTCAGATGACTTCTGTACTCGAGTACAGAAGTCATCTGAGGAGG |
| H-sh4-Mar-F-2 | CCGG CCAAGACCTTCTCTTCCAGAT CTCGAGATCTGGAAGAGAAGGTCTTGGTTTTTG |
| H-sh4-Mar-R-2 | AATTCAAAAA CCAAGACCTTCTCTTCCAGATCTCGAG ATCTGGAAGAGAAGGTCTTGG |
| H-shIL1RL1-F-1 | CCGG CGCAGGTGATTACACCTGTAA CTCGAGTTACAGGTGTAATCACCTGCGTTTTTG |
| H-shIL1RL1-R-1 | AATTCAAAAA CGCAGGTGATTACACCTGTAA CTCGAGTTACAGGTGTAATCACCTGCG |
| H-shDENND2A-F-1 | CCGGCCTAGTGCAGCCCTATTCTTTCTCGAGAAAGAATAGGGCTGCACTAGGTTTTTG |
| H-shDENND2A-R-1 | AATTCAAAAACCTAGTGCAGCCCTATTCTTTCTCGAGAAAGAATAGGGCTGCACTAGG |
| H-shDENND2A-F-2 | CCGG CCAATGAAGGAGAACCCTTATCTCGAGATAAGGGTTCTCCTTCATTGGTTTTTG |
| H-shDENND2A-R-2 | AATTCAAAAA CCAATGAAGGAGAACCCTTATCTCGAG ATAAGGGTTCTCCTTCATTGG |
| H-shPRR5L-F-1 | CCGGTATGCGATGCTGCCTTATTTCCTCGAGGAAATAAGGCAGCATCGCATATTTTTG |
| H-shPRR5L-R-1 | AATTCAAAAATATGCGATGCTGCCTTATTTCCTCGAGGAAATAAGGCAGCATCGCATA |
| H-shIL11-F-1 | CCGG TGCACAGCTGAGGGACAAATTCTCGAGAATTTGTCCCTCAGCTGTGCATTTTTG |
| H-shIL11-R-1 | AATTCAAAAA TGCACAGCTGAGGGACAAATTCTCGAGAATTTGTCCCTCAGCTGTGCA |
| H-shFLNC-F-1 | CCGGCGGTGTGTCATCAGAGTTCATCTCGAGATGAACTCTGATGACACACCGTTTTTG |
| H-shFLNC-R-1 | AATTCAAAAACGGTGTGTCATCAGAGTTCATCTCGAGATGAACTCTGATGACACACCG |

| | |
|---|---|
| H-shLAMC2-F-1 | CCGGGCCCTGCAATTGTAACTCCAACTCGAGTTGGAGTTACAATTGCAGGGCTTTTTG |
| H-shLAMC2-R-1 | AATTCAAAAAGCCCTGCAATTGTAACTCCAACTCGAGTTGGAGTTACAATTGCAGGGC |
| H-shHBEGF-F-1 | CCGG CTTCTCATGTTTAGGTACCATCTCGAG ATGGTACCTAAACATGAGAAGTTTTTG |
| H-shHBEGF-R-1 | AATTCAAAAA CTTCTCATGTTTAGGTACCAT CTCGAG ATGGTACCTAAACATGAGAAG |
| H-shCTGF-F-1 | CCGG CATCTTTGAATCGCTGTACTACTCGAGTAGTACAGCGATTCAAAGATG TTTTTG |
| H-shCTGF-R-1 | AATTCAAAAA CATCTTTGAATCGCTGTACTA CTCGAG TAGTACAGCGATTCAAAGATG |
| H-shMSN-F-1 | CCGGACCACCGGGAAGCAGCTATTTCTCGAGAAATAGCTGCTTCCCGGTGGTTTTTTG |
| H-shMSN-R-1 | AATTCAAAAAACCACCGGGAAGCAGCTATTTCTCGAGAAATAGCTGCTTCCCGGTGGT |
| H-shCYR61-F-1 | CCGGTTGAGGAGCATTAAGGTATTTCTCGAGAAATACCTTAATGCTCCTCAATTTTTG |
| H-shCYR61-R-1 | AATTCAAAAATTGAGGAGCATTAAGGTATTTCTCGAGAAATACCTTAATGCTCCTCAA |
| H-shTMEM171-F-1 | CCGGTGTAGGTGACTCGGTAATAATCTCGAGATTATTACCGAGTCACCTACATTTTTG |
| H-shTMEM171-R-1 | AATTCAAAAATGTAGGTGACTCGGTAATAATCTCGAGATTATTACCGAGTCACCTACA |
| H-shAXL-F-1 | CCGGCGAAAGAAGGAGACCCGTTATCTCGAGATAACGGGTCTCCTTCTTTCGTTTTTG |
| H-shAXL-R-1 | AATTCAAAAACGAAAGAAGGAGACCCGTTATCTCGAGATAACGGGTCTCCTTCTTTCG |
| H-shAXL-F-2 | CCGGGCTGTGAAGACGATGAAGATTCTCGAGAATCTTCATCGTCTTCACAGCTTTTTG |
| H-shAXL-R-2 | AATTCAAAAAGCTGTGAAGACGATGAAGATTCTCGAGAATCTTCATCGTCTTCACAGC |
| H-shNUP214-F-1 | CCGGCCTTTCGATCCCTGGGACCAACTCGAGTTGGTCCCAGGGATCGAAAGGTTTTTG |
| H-shNUP214-R-1 | AATTCAAAAACCTTTCGATCCCTGGGACCAACTCGAGTTGGTCCCAGGGATCGAAAGG |
| H-shEPB41L2-F-1 | CCGGTAGTGAACTCAAGCGCAATTTCTCGAGAAATTGCGCTTGAGTTCACTATTTTTG |
| H-shEPB41L2-R-1 | AATTCAAAAATAGTGAACTCAAGCGCAATTTCTCGAGAAATTGCGCTTGAGTTCACTA |
| H-shNCF2-F-1 | CCGGGCGCAACTACAGATTTGGAAACTCGAGTTTCCAAATCTGTAGTTGCGCTTTTTG |
| H-shNCF2-R-1 | AATTCAAAAAGCGCAACTACAGATTTGGAAACTCGAGTTTCCAAATCTGTAGTTGCGC |
| H-shRASGRF-F-1 | CCGGTGAGGATGATATCCCATATTACTCGAGTAATATGGGATATCATCCTCATTTTTG |
| H-shRASGRF-R-1 | AATTCAAAAATGAGGATGATATCCCATATTACTCGAGTAATATGGGATATCATCCTCA |
| H-shRASGRF-F-2 | CCGGGCCTCCTTATATTGTGATGATCTCGAGATCATCACAATATAAGGAGGCTTTTTG |

| | |
|---|---|
| H-shRASGRF-R-2 | AATTCAAAAAGCCTCCTTATATTGTGATGATCTCGAGATCATCACAATATAAGGAGGC |
| Q-hGAPDH-F | TCCCTCAAGATTGTCAGCAA |
| Q-hGAPDH-R | AGATCCACAACGGATACATT |
| Q-hAXL-F | CAGTGCCAAATCCGGGGAG |
| Q-hAXL-R | TGGGTGCCAAACTTTCCTCA |
| Q-hGCLC-F | GGCACAAGGACGTTCTCAAGT |
| Q-hGCLC-R | CAGACAGGACCAACCGGAC |
| Q-hGCLM-F | TGTCTTGGAATGCACTGTATCTC |
| Q-hGCLM-R | CCCAGTAAGGCTGTAAATGCTC |
| Q-hBCKDHA-F | CTACAAGAGCATGACACTGCTT |
| Q-hBCKDHA-R | CCCTCCTCACCATAGTTGGTC |
| Q-hBCKDK-F | CAAGACCGTCACCTCCTTTTAC |
| Q-hBCKDK-R | GGCCAGCGTAGAGCATCAT |
| Q-hBCAT1-F-1 | CCAAAGCCCTGCTCTTTGTA |
| Q-hBCAT1-R-1 | TGGAGGAGTTGCCAGTTCTT |
| Q-hBCAT1-F-2 | CCAAAGCCCTGCTCTTTGTA |
| Q-hBCAT1-R-2 | TGGAGGAGTTGCCAGTTCTT |
| Q-hBMP5-F | GCTGCTGGGTTCTAGTGGG |
| Q-hBMP5-R | TTCGTGGTTCCGTAGTCTTCTA |
| Q-hFAM171A1-F | GCAGATGCGCTCATCGAGAT |
| Q-hFAM171A1-R | GCCCAGCTTATACTGGAACTTG |
| Q-hCYB5R2-F | AGGAGGAGAGAGCCAATCACC |
| Q-hCYB5R2-R | ACATAGTTACCTACAGGAAGCCC |
| Q-hHSD3B1-F | CACATGGCCCGCTCCATAC |
| Q-hHSD3B1-R | GTGCCGCCGTTTTTCAGATTC |

| | |
|---|---|
| Q-h4-Mar-F | TACAAGTACCACGTCATCGC |
| Q-h4-Mar-R | AGTTGACCAGATGAGCCAAG |
| Q-hIL1RL1-F | GAAAACCTAGTTACACCGTGGAT |
| Q-hIL1RL1-R | GCAAACACACGATTTCTTTCCTG |
| Q-hDENND2A-F | AGCTCAGAGGTGTTCAGAACC |
| Q-hDENND2A-R | CAGATAATCCTCCTGTCCGTCT |
| Q-hPRR5L-F | CGGCTGTTGAAGAGTGAACTT |
| Q-hPRR5L-R | GCAGGGTAGGGAGAGTCTCAG |
| Q-hIL11-F | CGAGCGGACCTACTGTCCTA |
| Q-hIL11-R | GCCCAGTCAAGTGTCAGGTG |
| Q-hFLNC-F | CTGGGCGATGAGACAGACG |
| Q-hFLNC-R | GCGGATGGAACTTGCGGTA |
| Q-hLAMC2-F | GACAAACTGGTAATGGATTCCGC |
| Q-hLAMC2-R | TTCTCTGTGCCGGTAAAAGCC |
| Q-hBEGF-F | ATCGTGGGGCTTCTCATGTTT |
| Q-hBEGF-R | TTAGTCATGCCCAACTTCACTTT |
| Q-hCTGF-F | CAGCATGGACGTTCGTCTG |
| Q-hCTGF-R | AACCACGGTTTGGTCCTTGG |
| Q-hMSN-F | ATGCCCAAAACGATCAGTGTG |
| Q-hMSN-R | ACTTGGCACGGAACTTAAAGAG |
| Q-hCYR61-F | CTCGCCTTAGTCGTCACCC |
| Q-hCYR61-R | CGCCGAAGTTGCATTCCAG |
| Q-hTMEM171-F | GGACAGACACGTCAGCAAACT |
| Q-hTMEM171-R | GGGAGGGGCTTATATTGGCAT |
| Q-hAXL-F | CAGTGCCAAATCCGGGGAG |

| | |
|---|---|
| Q-hAXL-R | TGGGTGCCAAACTTTCCTCA |
| Q-hNUP214-F | TGACTCCCCTGAGGAATTGC |
| Q-hNUP214-R | GCGAAGACCAGACCATATTTGTT |
| Q-hEPB41L2-F | AGAATCAGTCTTCCGATCCAGA |
| Q-hEPB41L2-R | TGAACCGAGAAATACCCCTGC |
| Q-hNCF2-F | CCCACTCCCGGATTTGCTTC |
| Q-hNCF2-R | GTCTCGGTTAATGCTTCTGGTAA |
| Q-hRASGRF-F | GCCCTTTACAGTCATCCTAGTG |
| Q-hRASGRF-R | GCACAGTGACCTTGGAATTTTC |
| Q-hBCAT1-promoter-F | ATTTAGTAAGATACAGGGAAGACA |
| Q-hBCAT1-promoter-R | TGTTCCGTCGGCCACGAGGGAA |
| hBCAT1-CpG-F | AGTTTGAGAGTAGTTTGGGTAATATAGTAA |
| hBCAT1-CpG-R | AATATCCCTAAAAATCCAAAATC |
| Q-hMET amplification-F-1 | GTTCATGCCGACAAGTGCAG |
| Q-hMET amplification-R-1 | TGCACAATCAGGCTACTGGG |
| Q-hMET amplification-F-2 | CCATCCAGTGTCTCCAGAAGTG |
| Q-hMET amplification-R-2 | TTCCCAGTGATAACCAGTGTGTAG |
| Q-hHER2 amplification-F-1 | CATTGGGACCGGAGAAACCA |
| Q-hHER2 amplification-R-1 | TCCATGGTGCTCACTGCG |
| Q-hHER2 amplification-F-2 | GGTGGGGACCTGACACTAGG |
| Q-hHER2 amplification-R-2 | GTCATGTGTGGGGAGGCTTT |
| Q-hMTHFR amplification-F-1 | CCATCTTCCTGCTGCTGTAACTG |
| Q-hMTHFR amplification-R-1 | GCCTTCTCTGCCAACTGTCC |
| SUV39H1-F | ATGGTGGGGATGAGTCGCC |
| SUV39H1-R | CTAGAAGAGGTATTTGCGGCAG |

| | |
|---|---|
| Xho1-SUV39H1-F | CCGCTCGAGGCCACCATGGTGGGGATGAGTCGCC |
| SUV39H1-BAMH1-R | CGGGATCCCTAGAAGAGGTATTTGCGGCAG |
| Q-hSUV39H1-F | CAAGTTTGCCTACAATGACCAGG |
| Q-hSUV39H1-R | GTACCACACGATTTGGGCAGT |
| H-ShSUV39H1-F-1 | CCGGCGTTGGGATTCATGGCCTATTCTCGAG AATAGGCCATGAATCCCAACGTTTT TG |
| H-ShSUV39H1-R-1 | AATTCAAAAACGTTGGGATTCATGGCCTATTCTCGAG AATAGGCCATGAATCCCAACG |
| H-ShSUV39H1-F-2 | CCGGAGTCGAGTACCTGTGCGATTACTCGAG TAATCGCACAGGTACTCGACTTTTT TG |
| H-ShSUV39H1-R-2 | AATTCAAAAAAGTCGAGTACCTGTGCGATTACTCGAG TAATCGCACAGGTACTCGACT |
| G9a-F | ATGCGGGGTCTACCGAGA |
| G9a-R | TCATGTGTTGACAGGGGGCA |
| XBA-G9a-F | GCTCTAGAGCCACC ATGCGGGGTCTACCGAGA |
| G9a-NOT1-R | ATAGTTTAGCGGCCGC TCATGTGTTGACAGGGGGCA |
| Q-hG9a-F | GGGCGGGAAAATCACCTCC |
| Q-hG9a-R | CACTCATGCGGAAATGCTGTAT |
| H-ShG9a-F-1 | CCGGCGAGAGAGTTCATGGCTCTTTCTCGAG AAAGAGCCATGAACTCTCTCGTTTTTG |
| H-ShG9a-R-1 | AATTCAAAAACGAGAGAGTTCATGGCTCTTTCTCGAGAAAGAGCCATGAACTCTCTCG |
| H-ShG9a-F-2 | CCGGCTCCAGGAATTTAACAAGATTCTCGAGAATCTTGTTAAATTCCTGGAGTTTTTG |
| H-ShG9a-F-2 | AATTCAAAAACTCCAGGAATTTAACAAGATTCTCGAGAATCTTGTTAAATTCCTGGAG |
| Xho-G9a-F | CCGCTCGAGGCCACCATGGCGGCGGCGGCGGGAGCTGCA |
| G9a-Ecor-R | CGGAATTTCATGTGTTGACAGGGGGCA |
| BCAT1-F | CCAAAGCCCTGCTCTTTGTA |
| BCAT1-R | TGGAGGAGTTGCCAGTTCTT |
| XHO1-BCAT1-F | CCGctcgagGCCACCATGAAGGATTGCAGTAACGGA |
| BCAT1-NHE1-R | TTTTCCTTTTGCGGCCGCTCAGGATAGCACAATTGT |
| h-shALK-1-F | CCGGACCCAAATCAAGAAACCTGTTCTCGAGAACAGGTTTCTTGATTTGGGTTTTTTG |

| | |
|---|---|
| h-shALK-1-R | AATTCAAAAAACCCAAATCAAGAAACCTGTTCTCGAGAACAGGTTTCTTGATTTGGGT |
| h-shALK-2-F | CCGGGTGATAAATACAAGGCCCAGACTCGAGTCTGGGCCTTGTATTTATCACTTTTTG |
| h-shALK-2-R | AATTCAAAAAGTGATAAATACAAGGCCCAGACTCGAGTCTGGGCCTTGTATTTATCAC |
| Q-hHER2-F | GAGGGCCGGTATACATTCGG |
| Q-hHER2-R | CCCAGACCATAGCACACTCG |
| Q-hALK-F | AGAACTCCAACCTGAGCGTG |
| Q-hALK-R | CGTAGCCCGAGATGCTGTAG |
| sh-hJMJD1a-1-F | CCGGGCTTTGATTGTGAAGCATTTACTCGAGTAAATGCTTCACAATCAAAGCTTTTTG |
| sh-hJMJD1a-1-R | AATTCAAAAAGCTTTGATTGTGAAGCATTTACTCGAGTAAATGCTTCACAATCAAAGC |
| sh-hJMJD1a-2-F | CCGGACAACTTAGATTGGGTTTATACTCGAGTATAAACCCAATCTAAGTTGTTTTTTG |
| sh-hJMJD1a-2-R | AATTCAAAAAACAACTTAGATTGGGTTTATACTCGAGTATAAACCCAATCTAAGTTGT |
| sh-hJMJD1b-1-F | CCGGCCCTAGTTCATCGCAACCTTTCTCGAGAAAGGTTGCGATGAACTAGGGTTTTTG |
| sh-hJMJD1b-1-R | AATTCAAAAACCCTAGTTCATCGCAACCTTTCTCGAGAAAGGTTGCGATGAACTAGGG |
| sh-hJMJD1b-2-F | CCGGCACATAACTGGTACAAGTATGCTCGAGCATACTTGTACCAGTTATGTGTTTTTG |
| sh-hJMJD1b-2-R | AATTCAAAAACACATAACTGGTACAAGTATGCTCGAGCATACTTGTACCAGTTATGTG |
| sh-hJMJD2a-1-F | CCGGTTCGAGAGTTCCGCAAGATAGCTCGAGCTATCTTGCGGAACTCTCGAATTTTTG |
| sh-hJMJD2a-1-R | AATTCAAAAATTCGAGAGTTCCGCAAGATAGCTCGAGCTATCTTGCGGAACTCTCGAA |
| sh-hJMJD2a-2-F | CCGGTAGTGAAAGGACGAGCCATTTCTCGAGAAATGGCTCGTCCTTTCACTATTTTTG |
| sh-hJMJD2a-2-R | AATTCAAAAATAGTGAAAGGACGAGCCATTTCTCGAGAAATGGCTCGTCCTTTCACTA |
| sh-hJMJD2c-1-F | CCGGATACTTGGATTACGAAGATTTCTCGAGAAATCTTCGTAATCCAAGTATTTTTTG |
| sh-hJMJD2c-1-R | AATTCAAAAAATACTTGGATTACGAAGATTTCTCGAGAAATCTTCGTAATCCAAGTAT |
| sh-hJMJD2c-2-F | CCGGGCCCAAGTCTTGGTATGCTATCTCGAGATAGCATACCAAGACTTGGGCTTTTTG |
| sh-hJMJD2c-2-R | AATTCAAAAAGCCCAAGTCTTGGTATGCTATCTCGAGATAGCATACCAAGACTTGGGC |
| sh-hJMJD2d-1-F | CCGGCCTGGAATGTCTTTGGATATTCTCGAGAATATCCAAAGACATTCCAGGTTTTTG |
| sh-hJMJD2d-1-R | AATTCAAAAACCTGGAATGTCTTTGGATATTCTCGAGAATATCCAAAGACATTCCAGG |

| | |
|---|---|
| sh-hJMJD2d-2-F | CCGGCGCATCTATAATTCACCGATTCTCGAGAATCGGTGAATTATAGATGCGTTTTTG |
| sh-hJMJD2d-2-R | AATTCAAAAACGCATCTATAATTCACCGATTCTCGAGAATCGGTGAATTATAGATGCG |
| sh-hJMJD1c-1-F | CCGGGCTCCTGTGATTCAATGTTATCTCGAGATAACATTGAATCACAGGAGCTTTTTG |
| sh-hJMJD1c-1-R | AATTCAAAAAGCTCCTGTGATTCAATGTTATCTCGAGATAACATTGAATCACAGGAGC |
| sh-hJMJD1c-2-F | CCGGCCCATTTACTAGCCGGATCATCTCGAGATGATCCGGCTAGTAAATGGGTTTTTG |
| sh-hJMJD1c-2-R | AATTCAAAAACCCATTTACTAGCCGGATCATCTCGAGATGATCCGGCTAGTAAATGGG |
| sh-hJMJD2b-1-F | CCGGGCGGCAGACGTATGATGACATCTCGAGATGTCATCATACGTCTGCCGCTTTTTG |
| sh-hJMJD2b-1-R | AATTCAAAAAGCGGCAGACGTATGATGACATCTCGAGATGTCATCATACGTCTGCCGC |
| sh-hJMJD2b-2-F | CCGGGATGACCTTGAACGCAAATACCTCGAGGTATTTGCGTTCAAGGTCATCTTTTTG |
| sh-hJMJD2b-2-R | AATTCAAAAAGATGACCTTGAACGCAAATACCTCGAGGTATTTGCGTTCAAGGTCATC |
| Q-hJMJD1C-F | CAGGTCTCGTGCCAATCAAAA |
| Q-hJMJD1C-R | GCTGTTGCTGGTGTGTATTCT |
| Q-hJMJD1A-F | GTGCTCACGCTCGGAGAAA |
| Q-hJMJD1A-R | GTGGGAAACAGCTCGAATGGT |
| Q-hJMJD1B-F | CCCCAGGACCTAGCGATCTT |
| Q-hJMJD1B-R | AGCAGAAGCACGATAACTTCC |
| Q-hJMJD2A-F | ATCCCAGTGCTAGGATAATGACC |
| Q-hJMJD2A-R | ACTCTTTTGGAGGAACAACCTTG |
| Q-hJMJD2B-F | ACTTCAACAAATACGTGGCCTAC |
| Q-hJMJD2B-R | CGATGTCATCATACGTCTGCC |
| Q-hJMJD2D-F | GGGCAGGGGTGTTTACTCAAT |
| Q-hJMJD2D-R | TGTTTGCCAAATGGCGATACT |
| Q-hJMJD2C-F | CGAGGTGGAAAGTCCTCTGAA |
| Q-hJMJD2C-R | GGGCTCCTTTAGACTCCATGTAT |
